# From Ligand Binding to Pore Opening in the Mosquito Odorant Receptor AaegOR10

**DOI:** 10.64898/2026.09.28.754911

**Authors:** Tiefeng Song, Zhanyi Xu, Zheyi Liu, Mengli Chen, Yan Ru, Yirong Guo, Yong Wang

## Abstract

Mosquito olfaction is central to host seeking and other behaviors relevant to vector ecology and disease transmission. Mosquito odorant receptors (ORs) are ligand-gated ion channels, yet how ligand binding is coupled to pore opening remains poorly understood. Here, we combine more than 400 *µ*s of aggregate molecular dynamics simulations and adaptive sampling with Markov state models (MSMs) and electrophysiological measurements to characterize the gating dynamics of *Aedes aegypti* odorant receptor 10 (AaegOR10). Interaction analysis identifies M288 as a persistent contact between *o*-cresol and S6, connecting the orthosteric pocket to a conformation-dependent hydrophobic network involving I293, F294, and F356 at the S6-S7b interface. The MSMs place the experimentally determined open and closed structures within a multistate conformational landscape and identify a principal intermediate along the dominant gating pathway. A localized S6 kink in this state is associated with weakened S6-S7b coupling and increased flexibility of the pore-lining S7b helix. Ligand binding shifts the conformational equilibrium toward open states and shortens the transition times from the closed state to both the intermediate and open states. We also identify an off-path closed state characterized by a pronounced inward displacement of S6, with phospholipid headgroup insertion observed in some sampled conformations. Mutational analysis and voltage-clamp recordings support the functional relevance of M288, the S6-S7b interface, and the kink-prone S6 region. Together, these results describe a multistate pathway linking ligand recognition to AaegOR10 pore gating.

## Introduction

Globally distributed mosquitoes transmit several major human pathogens, while their preference for human hosts makes odor-guided behavior directly relevant to public health.^1–4^ Odor cues shape mosquito behaviors, including seeking hosts, locating food sources, finding mating partners, and selecting oviposition sites. For disease-transmitting mosquitoes, odor-guided host seeking is also an important component of vector behavior and therefore has direct relevance to disease transmission and vector control.^5–11^ Understanding how mosquito olfactory systems recognize chemical cues and convert them into neuronal signals is thus important both for defining the molecular basis of insect sensory behavior and for identifying potential targets for disrupting ecologically important behaviors.

Insect odorant receptors (ORs) are central components of the insect olfactory system. Unlike mammalian ORs, which belong to the G-protein-coupled receptor (GPCR) family, insect ORs form ligand-gated, non-selective cation channels that directly couple odorant binding to membrane depolarization.^12^ Most insect ORs function as heteromeric complexes composed of a divergent OR subunit, which determines ligand specificity, and the more conserved odorant receptor co-receptor, Orco.^13,14^ The transmembrane helices of the OR subunit form a hydrophobic pocket capable of recognizing chemically diverse ligands, ^15,16^ whereas Orco contributes to receptor assembly, membrane trafficking, and ion conduction.^17,18^ Both subunits contain seven transmembrane helices, but with an inverted topology relative to GPCRs, placing the N-terminus intracellularly and the C-terminus extracellularly.^19^

Recent cryo-electron microscopy (cryo-EM) structures have provided an atomic-level view of ligand recognition and pore gating in insect ORs, including the mosquito receptor *Aedes aegypti* AaegOR10, *Anopheles gambiae* AgamOR28, *D. melanogaster* DmOR67d and *D. bipectinata* DbOR67d and the pea aphid *Acyrthosiphon pisum* ApisOR5 and ApisOR22.^20–23^ These structures establish a common 1:3 OR stoichiometry, in which one OR subunit is associated with three Orco subunits to form the functional channel. AaegOR10 from *Aedes aegypti* responds to several odorants, including *o*-cresol, which serves as an activating ligand for the receptor.^20^ In AaegOR10, the principal ligand-binding pocket is formed by S2, S3, S4, and S6, whereas pore opening involves the S5-S7 region. Comparison of the ligand-free closed and ligand-bound open structures further suggests that the ligand-binding region is relatively rigid, whereas the S5-S7 region undergoes the major conformational rearrangements associated with channel opening. In particular, displacement of the pore-lining S7b helix converts a narrow hydrophobic constriction into a wider conducting pathway.

These structures define experimentally accessible conformational endpoints, but they do not resolve the dynamic pathway connecting them. ^24,25^ A central question is therefore how ligand binding within the relatively rigid S1-S4 region is mechanically coupled to rearrangement of the more mobile S5-S7 gating module. The interactions that transmit this signal, the conformational intermediates along the gating pathway, and the kinetic barriers separating these states remain unclear. Moreover, the membrane environment may contribute directly to receptor dynamics. Unassigned densities observed near the transmembrane helices in cryo-EM apo structures suggest possible lipid interactions, but the implications of these observations remain unclear.

Here, we combine extensive molecular dynamics (MD) simulations, adaptive sampling with 400+ *µ*s of aggregate simulation data, and Markov state model (MSM) analysis to characterize the conformational and kinetic landscape of AaegOR10 gating.^26,27^ Using the cryo-EM open and closed structures as experimentally defined reference states, we resolve the intermediate states and conformational pathways connecting them. This analysis identifies M288 as a key link between the ligand-binding pocket and the S6-S7b gating interface and reveals a conformational mechanism in which rearrangement of the S6-S7b hydrophobic network accompanies progression toward pore opening. Ligand binding shifts the conformational equilibrium toward open states and shortens the transition times from the closed state to both the principal intermediate and open states. We further identify an off-path closed state characterized by a pronounced inward displacement of S6. Targeted mutations and voltage-clamp measurements provide experimental tests of the residue-level interactions identified by the simulations. By delineating the allosteric pathway from ligand binding to pore opening, this study provides a structural framework for designing inhibitors beyond those targeting the orthosteric pocket alone.

## Results

### M288 Couples Ligand Binding to Channel Opening

The cryo-EM structures of apo-closed (AaegOR10_Apo-Closed_) and *o*-cresol-bound open (AaegOR10_Holo-Open_) AaegOR10 provide two structural endpoints for the gating transition. Comparison of the two structures shows that the S1-S4 bundle is comparatively rigid, whereas the largest conformational rearrangements occur within the S5-S7 region (Fig. 1a). To examine how ligand binding is coupled to these conformational changes, we performed 3 *µ*s of unbiased MD for each of four systems: the two experimentally determined states and two cross-state systems, consisting of ligand-bound closed (AaegOR10_Holo-Closed_) and apo open (AaegOR10_Apo-Open_) receptors. This four-system setup enables differentiation of ligand-dependent molecular interactions from conformational changes driven solely by receptor structural states.

**Figure 1:**
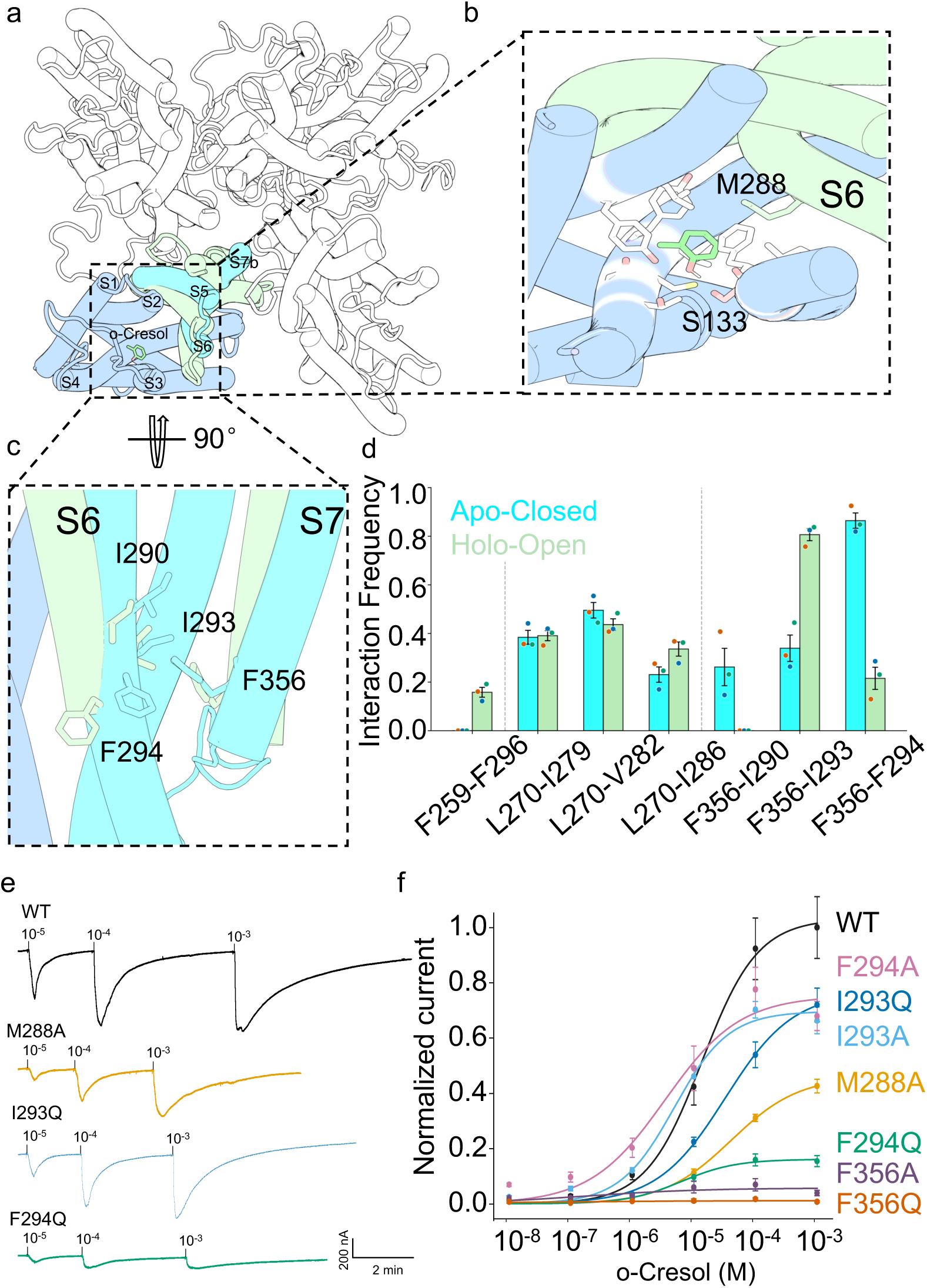
Ligand coupling and mutational effects. a) Apo-closed and holo-open cryo-EM structures showing S5-S7b rearrangements. b) Representative *o*-cresol binding pose with S133 and M288. c) S6-S7b hydrophobic interface. d) Hydrophobic interaction frequencies. Bars show means, error bars SE, and points individual trajectory values (*n* = 3). e) Representative current traces of WT AaegOR10, M288A, I293Q, and F294Q at the indicated *o*-cresol concentrations. Scale bars, 200 nA and 2 min. f) Normalized *o*-cresol concentration-response curves for WT AaegOR10 and the indicated mutants. Currents were normalized to the WT mean maximum response at 10*^−^*^3^ M. Points show means, error bars SE, and lines Hill fits; *n* = 5 or 6 independent recordings per construct.

Throughout the ligand-bound simulations, *o*-cresol remained well localized within the orthosteric pocket (Figs. 1b and S1a). Its hydroxyl group was oriented toward the S3-S4 cleft and formed a persistent hydrogen bond with S133, while the aromatic ring remained buried within the hydrophobic pocket. The distance between the ligand hydroxyl group and S133 showed a narrow distribution with little dependence on receptor conformation (Fig. S1b), indicating that this interaction provides a stable anchor for the ligand.

The interaction pattern between *o*-cresol and the receptor further distinguishes the ligand-binding region from the gating machinery. Across the 3 *µ*s trajectories, the ligand formed numerous contacts with the relatively rigid S1-S4 bundle, but only one persistent direct contact with the mobile S5-S7 module: the interaction between the ligand and M288 on S6 (Fig. S1d). Whereas the contacts within S1-S4 were largely preserved between conformational states, the M288 interaction varied with receptor conformation. Importantly, however, ligand-M288 contact was maintained even in the closed state, providing a continuous physical connection between the bound ligand and the S6 helix before pore opening.

To determine whether M288 is part of a broader communication pathway, we analyzed the interaction network connecting the ligand-binding pocket to the distal gating region using MDPath.^28^ A high-flux pathway propagated from the ligand-binding pocket toward the S5-S7 region through M288, whereas the alternative I132-A283 pathway remained largely confined to S6 (Fig. S2b). Taken together, both local interaction and network analyses demonstrate that M288 serves as a coupling site bridging the ligand-binding pocket and the dynamic gating module.

We next tested whether this simulated ligand-S6 interaction has a measurable effect on receptor function. M288 was replaced with alanine to reduce side-chain volume while retaining a hydrophobic character. Voltage-clamp recordings showed that M288A reduced the response to *o*-cresol relative to WT AaegOR10 (Figs. 1e, 1f, S4, and S5). The functional effect is consistent with a role for M288 in coupling the ligand-binding pocket to the gating machinery.

### A Hydrophobic Switch at the S6-S7b Interface Encodes Gating States

Having identified M288 as a connection between the ligand-binding pocket and S6, we next asked how the signal is transmitted from S6 to the pore-lining S7b helix. The MDPath network places the distal S6-S7b region downstream of M288 (Fig. S2b), suggesting that interactions within this interface may provide the next step in the gating pathway. We therefore examined hydrophobic contacts among S5, S6, and S7b across the four simulated conformational states.

The S6-S7b interface contains a cluster of hydrophobic residues, including I290, I293, F294, and F356. Its packing pattern changes markedly between the apo-closed and holo-open states (Fig. 1c,d). In the closed state, F356 forms distinct pairwise contacts with I293 and F294, and these relationships are substantially rearranged in the open state. The same state-dependent relationship is observed in the holo-closed and apo-open systems (Fig. S3a), indicating that the interaction pattern primarily reflects receptor conformation rather than ligand occupancy alone. In the open state, lateral displacement of S6 increases the separation between S6 and S7b and is accompanied by reduced I290-F356 interaction.

The F294-F356 pair is particularly sensitive to this rearrangement. These residues exhibit a strong co-evolutionary score (0.95), and their distance distribution changes from a relatively compact population to a bimodal distribution in the open ensemble, with a distinct long-distance population (Fig. S3b-d). In contrast, the I290-F356 and I293-F356 distances remain predominantly unimodal and compact. These observations identify F294-F356 as a prominent conformational switch within the S6-S7b interface.

The S5-S6 interface behaves differently. The main hydrophobic contacts involving I279, V282, I286, and L270 remain present across the simulated states, whereas the F296-F259 interaction is more transient (Fig. S3). Thus, the two interfaces do not undergo equivalent rearrangements during gating: the S5-S6 interface provides a relatively persistent structural anchor, while the S6-S7b interface undergoes state-dependent repacking.

To test the functional relevance of this interface, we introduced A and Q substitutions at I293, F294, and F356 to reduce side-chain volume or introduce hydrophilicity, respectively. Voltage-clamp recordings revealed residue-dependent changes in the concentration-response relationships to *o*-cresol (Figs. 1e, 1f, S4, and S5). The I293 substitutions produced intermediate effects. F294Q strongly suppressed the response, while F294A retained a substantial maximal response and produced a higher response than WT at low ligand concentrations. The enhanced response of F294A at low ligand concentrations is consistent with reduced stability of the closed state. At high ligand concentrations, the reduced maximal responses of F294A and F294Q indicate that F294 also contributes to open-state stability and overall receptor function. The F356 substitutions caused the strongest reduction among the mutations targeting the S6-S7b interface.

These results support a model in which ligand binding is transmitted through M288 to S6 and subsequently converted into a conformational rearrangement of the S6-S7b hydrophobic interface. The reciprocal packing of F356 with I293 and F294 provides a structural correlate of the closed-to-open transition.

### Thermodynamic Landscape of Allosteric Gating Revealed by MSM

The unbiased simulations resolved local interactions within the gating machinery but provided limited sampling of the complete transitions between the experimentally captured open and closed states. We therefore combined adaptive and unbiased simulations to characterize the broader conformational landscape. The resulting data comprised 276 *µ*s for the ligand-bound ensemble (270 *µ*s adaptive sampling plus 6 *µ*s unbiased MD) and 142 *µ*s for the apo ensemble (136 *µ*s adaptive sampling plus 6 *µ*s unbiased MD). Markov state models were constructed from these trajectories to quantify the populations and kinetics of the major conformational states (Fig. S6).

Projection of the conformational ensembles onto the first two time-lagged independent components (tIC1 and tIC2) produced a V-shaped free-energy surface (FES) (Fig. 2a). The apo trajectories were concentrated near the central vertex, corresponding to the closed region. The resting closed ensemble also displayed an elongated tail extending approximately perpendicular to the left arm of the V, indicating substantial fluctuations within the closed basin that are distinct from the principal open-closed transition.

**Figure 2:**
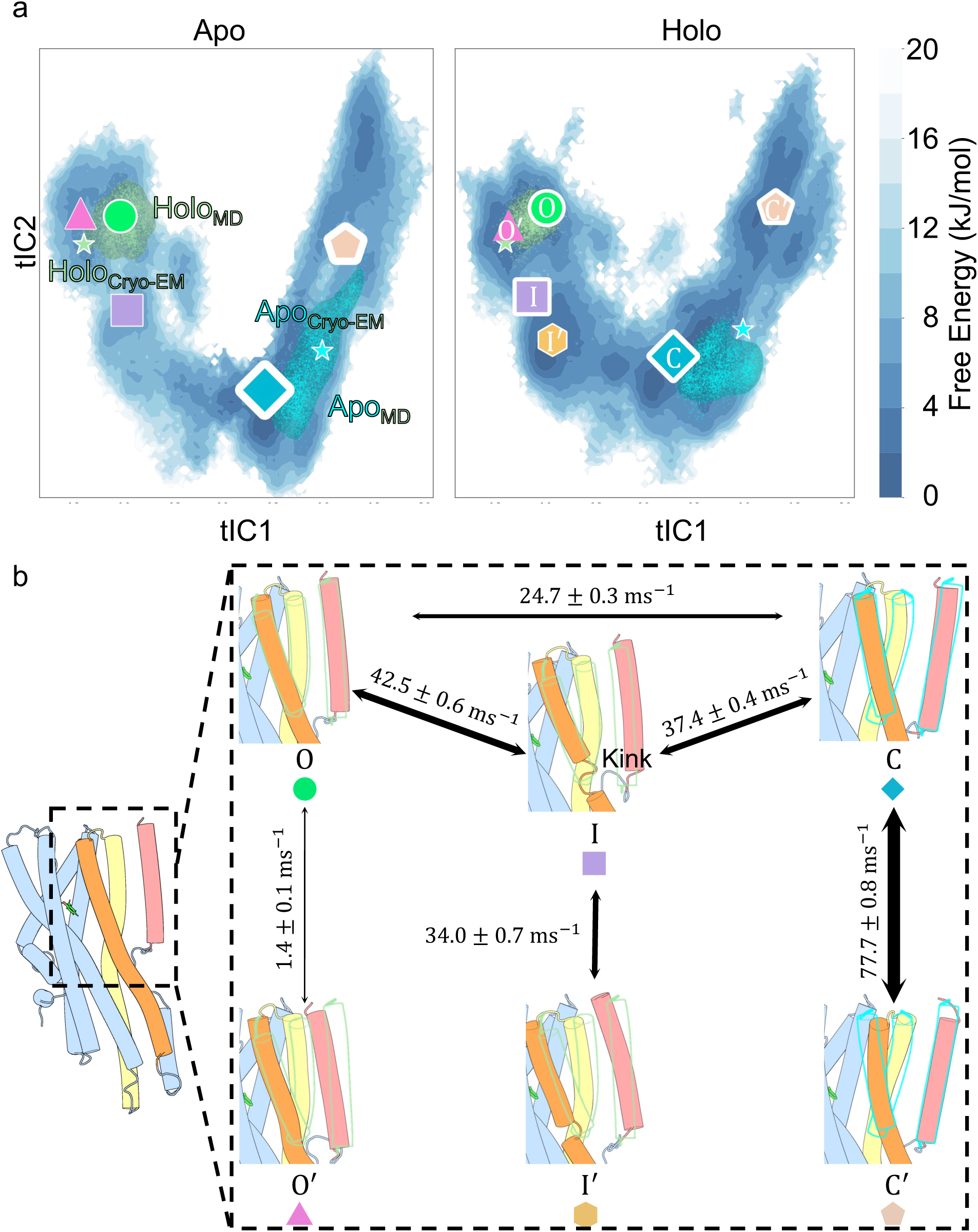
MSM free-energy surfaces and representative conformations. a) Apo and holo ensembles projected onto tIC1 and tIC2. Colored symbols denote macrostates, with white-outline size proportional to population. Light-blue and light-green points and shading show the apo and holo cryo-EM sampling ranges; stars mark the corresponding structures. b) Representative holo macrostates. Holo-open and apo-closed cryo-EM structures are outlined in light green and light blue, respectively. A1r0row widths and labels indicate pairwise total reactive fluxes and maximum-likelihood MSM values, with SE estimated from 100 transition-matrix samples in the holo BMSM ensemble.

The Koopman-reweighted FESs projected onto the tICA space and the minimum free-energy paths (MFEPs) further revealed four major conformational regions shared by the apo and ligand-bound ensembles: an open region at the left terminus of the V, a closed region at the central vertex, an intermediate region connecting them, and an off-path closed region along the upper-right branch (Figs. 2a and S7). The open region could be further separated into open and super-open subbasins. These regions provided a structural framework for examining how ligand binding changed both the equilibrium distribution and the accessibility of the gating pathway.

Comparison of the two FESs showed that *o*-cresol reshaped the conformational landscape. The ligand increased the population of open and intermediate conformations and lowered the barriers separating the major states.

### Macrostate Assignment and Kinetic Analysis of Ligand-Modulated Transitions

To quantify the kinetics associated with this landscape, the MSM microstates were coarse-grained using PCCA+. Five macrostates were identified in the apo ensemble and six in the ligand-bound ensemble (Figs. 2a and S8).

Five metastable states were common to both ensembles: open (O), intermediate (I), closed (C), super-open (O*^′^*), and off-path closed (C*^′^*). The simulated O and C macrostates were structurally consistent with the corresponding cryo-EM structures (Fig. 2b). The O*^′^* state exhibited pronounced outward displacements of S6 and S7b but constituted an exceedingly small fraction of both ensembles. The ligand-bound ensemble contained one additional off-path intermediate (I*^′^*) adjacent to I (Fig. 2).

MFPT calculations placed the global activation and deactivation processes on the millisecond timescale (Fig. S9a). Ligand binding preferentially accelerated activation-associated transitions from C, shortening the MFPTs from C to I and O, without uniformly accelerating all macrostate transitions. The kinetic effect of *o*-cresol is therefore state dependent and complements its redistribution of the conformational equilibrium toward open states.

Pairwise TPT calculations quantified the total reactive flux between each macrostate pair (Figs. 2b, S8, and S9b). Source-to-target pathway decompositions in both opening and closing directions identified routes through I as the major pathways in both ensembles, while direct O ⇌ C pathways remained accessible, particularly in the holo ensemble (Tables S3-S6).

### Structural Characteristics of the Intermediate State

The kinetic analysis identifies I as the dominant route between O and C. We next examined its structural basis and compared it with I*^′^*, which is unique to the ligand-bound ensemble.

Both I and I*^′^* retain an S5-S6 arrangement resembling that of O but contain a localized kink within the S291-H299 segment of S6 (Figs. 3a,b and S10a). The kink displaces I293 and F294 and weakens their hydrophobic interactions with S7b. Despite this common S6 configuration, I and I*^′^* differ in the behavior of S7b and in the resulting pore geometry. In I state, S7b remains highly dynamic, spanning a broad range between those of C and O. In I*^′^* state, by contrast, S7b remains predominantly open, and the minimum pore-distance distribution closely resembles that of O (Fig. S10b). Thus, I and I*^′^* share weakened S6-S7b coupling but differ in the extent to which S7b is stabilized in an open configuration.

**Figure 3:**
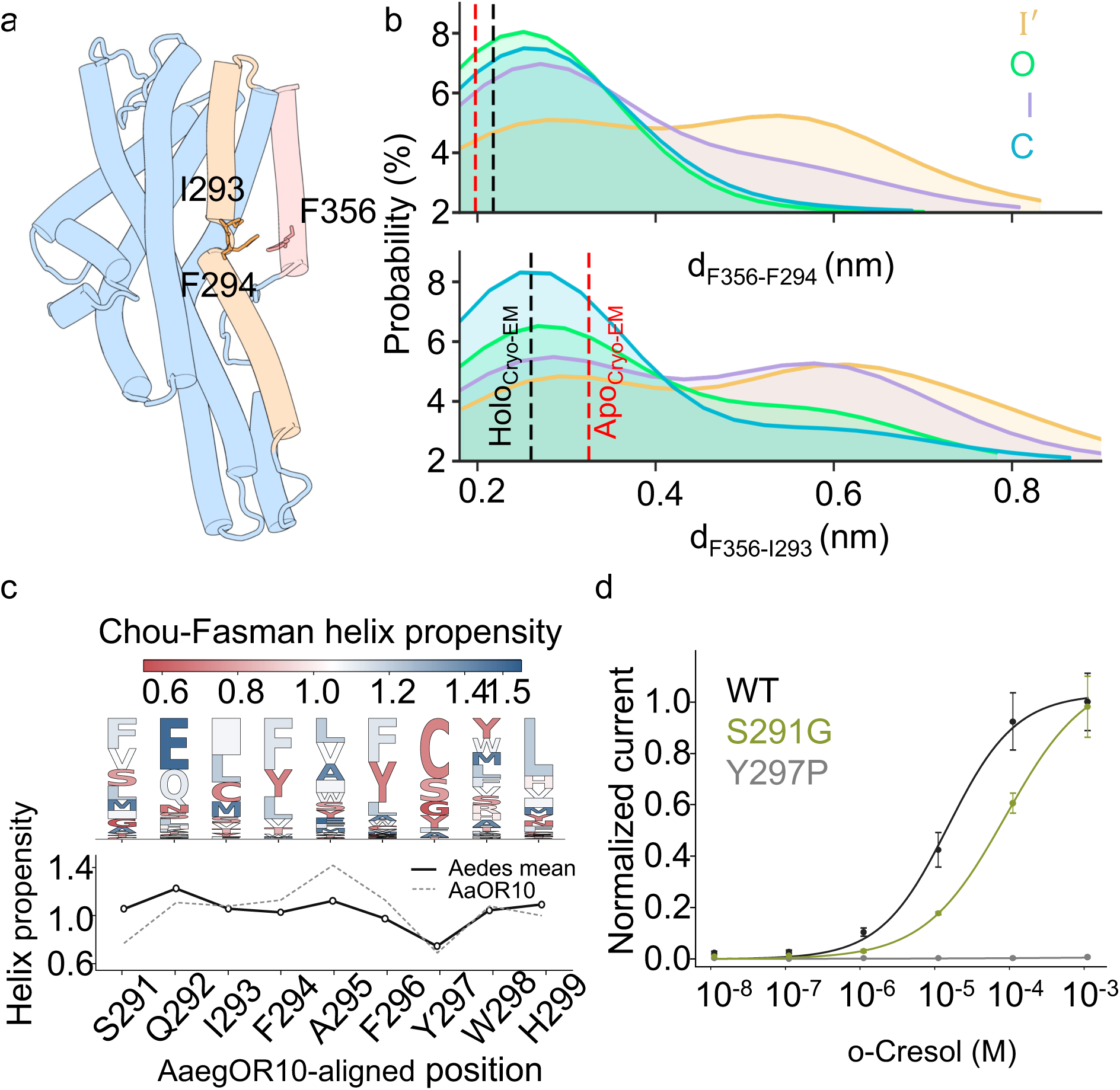
Structural characteristics of I and I*^′^*, helix propensity, and kink-region mutations. a) S291-H299 segment in I and I*^′^*. b) Distance distributions for the F356-I293 and F356-F294 pairs. c) FoldMason alignment and Chou-Fasman *α*-helix propensity analysis of the S291-H299 region across 176 *Aedes* OR-Orco systems. d) Normalized concentration-response relationships of WT AaegOR10, S291G, and Y297P. Points show means, error bars indicate SE, and lines show Hill fits; *n* = 5 or 6 independent recordings per construct.

**Figure 4:**
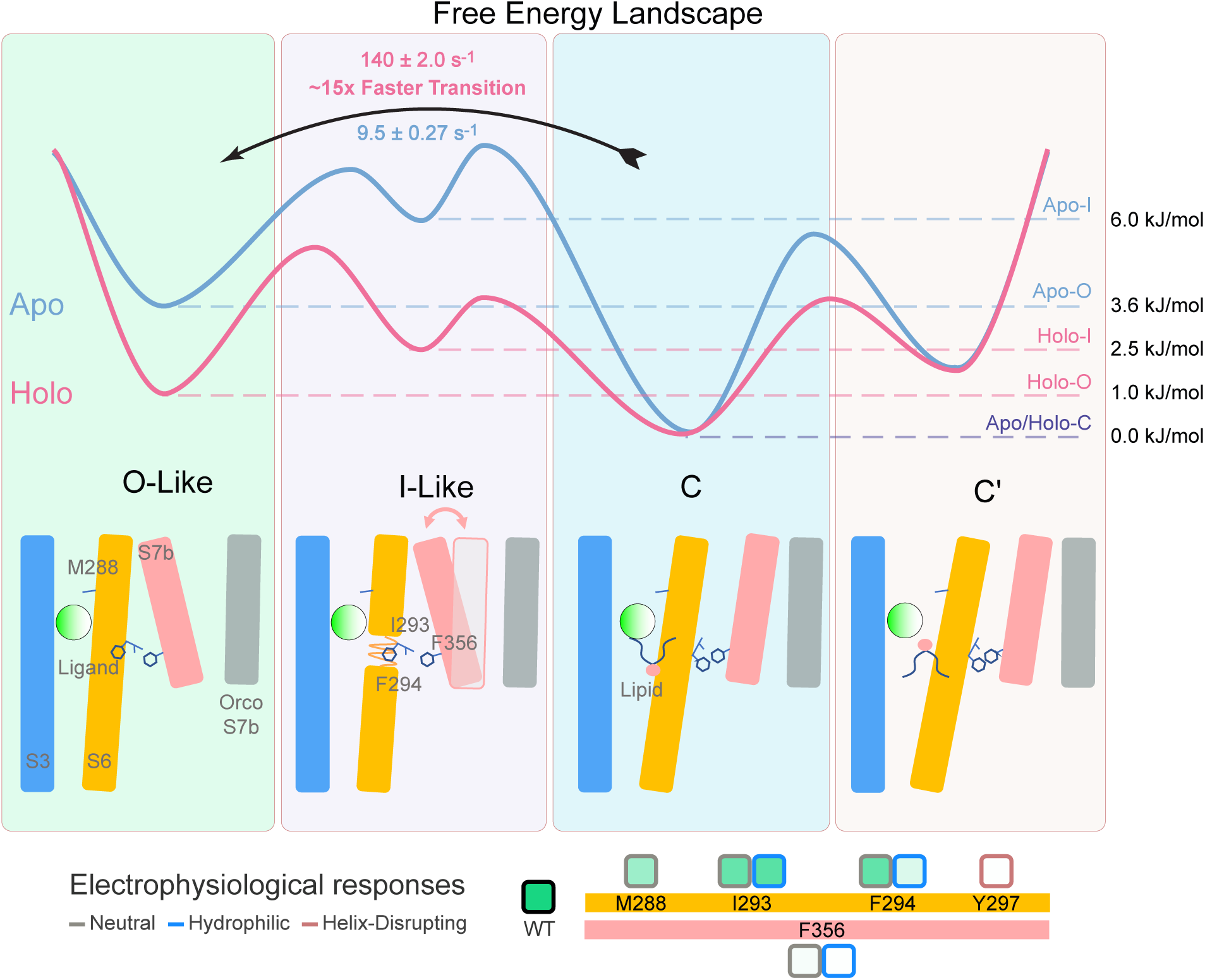
Summary of AaegOR10 conformational states and functional perturbations. The schematic integrates simulated free-energy profiles, representative structural features, state relationships, and normalized electrophysiological responses of WT AaegOR10 and the indicated mutants.

The macrostate populations further showed that *o*-cresol shifted the conformational landscape toward O and I while reducing the population of C*^′^*. Conformations assigned to O, O*^′^*, I*^′^*, and the open subpopulation of I represented more than 40% of the ligand-bound ensemble but less than 20% of the apo ensemble (Table S2). Thus, ligand binding does not simply stabilize a single open structure; it redistributes the receptor among several conformational states along and around the principal gating pathway.

The presence of the kink made us speculate that local secondary-structure properties of this S6 segment may influence gating. We therefore examined sequence and structural conservation across *Aedes* ORs. OR sequences were retrieved from VectorBase,^29^ modeled with AlphaFold3, ^30^ and structurally filtered to obtain 176 OR-Orco complexes, including 46 *Aedes aegypti* systems, 50 *Aedes albopictus* C6/36 systems, and 80 *Aedes albopictus* Foshan/FPA systems. The complexes were aligned with FoldMason,^31^ focusing on the region corresponding to S291-H299 of AaegOR10. Chou-Fasman analysis of the aligned sequences identified Y297 as having a pronounced helix-disrupting propensity (Fig. 3c). Together with the polar S291 residue within the kinked region, this result suggested that local destabilization of the S6 helix may be an intrinsic feature of the receptor.

We tested this hypothesis by introducing S291G and Y297P substitutions to perturb the local helical structure. The two mutations produced distinct functional phenotypes (Figs. 3d, S4, and S5). S291G reduced the response at subsaturating ligand concentrations but approached a maximal response similar to WT at saturation. This behavior is consistent with a change in the accessibility or stability of intermediate gating states without a major loss of the maximal conducting state. Y297P, in contrast, strongly reduced the response across the concentration range, indicating a more substantial disruption of receptor function.

Thus, the S291-H299 segment represents a structurally dynamic region within S6 that is coupled to the S6-S7b interface and influences the progression of the receptor through ligand-dependent gating states.

### Lipid Insertion Is Associated with Closed State

During the unbiased simulations, a POPC molecule entered the cleft between S3 and S6 from the inner leaflet in approximately half of the trajectories (Figs. S11a and S12). The location of the inserted lipid is compatible with the unassigned density observed near this region in the apo cryo-EM structure.

The conformational landscape also revealed a distinct off-path closed state. In C*^′^*, S5 and S7 remain relatively unchanged, whereas the upper portion of S6 shifts further inward relative to the apo-closed cryo-EM structure (Fig. S11b). A phospholipid headgroup was present in the S3-S6 cleft in some sampled C*^′^* conformations, with its acyl chain displaced and local hydration observed in the cleft (Fig. S11a). The C*^′^* population decreased from approximately 40% in the apo ensemble to approximately 23% in the ligand-bound ensemble (Fig. 2a), paralleling the ligand-induced redistribution toward intermediate and open conformations.

## Discussion and Conclusions

Structural studies of insect odorant receptors have established ligand-bound open and apo-closed conformations, but the pathway connecting these endpoints has remained unresolved.^20^ By combining molecular dynamics simulations, MSM analysis, and functional measurements, we place these structures within a multistate conformational landscape and identify interactions connecting the orthosteric pocket to the pore. The results indicate that AaegOR10 gating involves ligand-S6 coupling through M288, conformation-dependent rearrangement of the S6-S7b interface, and transitions among multiple metastable states (Fig. 4). Although ligand binding and release were not explicitly sampled here, a separate study of homomeric MhOR5 examined solvent-facing and membrane-facing binding/release pathways for ligands and quantified their association and dissociation kinetics.^32^

Within the AaegOR10 gating landscape resolved here, M288 occupies a distinctive position in the interaction network. Most ligand-contacting residues are located within the comparatively rigid S1-S4 bundle, whereas M288 provides a persistent contact between the ligand and the mobile S6 helix. The principal contacts at the S5-S6 interface remain comparatively stable across the simulated states, whereas the packing of the S6-S7b interface changes with receptor conformation. The mutational data support the functional relevance of these residues and interfaces: M288A reduces receptor responses, while substitutions at the S6-S7b interface produce residue- and concentration-dependent effects. In particular, the behavior of F294A and F294Q indicates that the contribution of F294 is not restricted to the closed state but also extends to open-state function and overall receptor activity.

The intermediate states clarify how similar local changes in S6 can be associated with different pore conformations. Both the principal intermediate and the off-path intermediate contain a localized S6 kink and weakened interactions between I293/F294 and F356, yet they differ in the behavior of S7b. S7b samples a broad range of pore conformations in the principal intermediate but remains predominantly open in the off-path intermediate. The S6 kink is therefore associated with increased S7b conformational flexibility but does not uniquely determine the pore conformation. Reactive-flux analysis places the principal intermediate on the main transition pathway between the open and closed states, although direct transitions between these states remain accessible. Ligand binding shifts the conformational equilibrium toward open states and shortens the transition times from the closed state to both the principal intermediate and open states. The distinct effects of S291G and Y297P further support the functional importance of the local helical properties of this S6 region.

The simulations also reveal a lipid-associated branch of the conformational landscape. In approximately half of the apo-closed unbiased trajectories, a POPC acyl chain entered the S3-S6 cleft from the inner leaflet. The inserted lipid is spatially compatible with the unassigned density in the apo cryo-EM structure, which may represent either a phospholipid headgroup or a sterol, as suggested by recent cryo-EM evidence.^33^ The off-path closed state showed a further inward position of the upper S6 helix relative to the apo-closed cryo-EM structure. Some sampled off-path closed conformations also contained a phospholipid headgroup in the S3-S6 cleft, together with local hydration.

Several limitations should be considered. Although the cumulative sampling provides access to transitions that are difficult to capture from cryo-EM structures or to resolve with existing AI-based generative approaches, additional metastable states or alternative transition pathways may remain unresolved. This possibility is supported by recent high-speed atomic force microscopy (HS-AFM) measurements showing that insect ORs undergo substantial conformational dynamics beyond the canonical tetrameric arrangement, including reversible square-to-pinched transitions and multiple kinetic substates. ^34^ These observations provide independent experimental evidence that insect ORs occupy a dynamic conformational landscape that extends beyond the static structures captured by cryo-EM. The conformational states resolved by HS-AFM and those identified here, however, describe different aspects of receptor dynamics. In addition, the correspondence between MSM-defined conformational states and functional channel states is not necessarily one-to-one. The principal intermediate identified here is kinetically connected to the open and closed states, but its conductance properties cannot be inferred directly from structural coordinates alone. Electrophysiological measurements provide functional support for the identified residues, but further experiments will be required to determine how individual conformational states map onto channel conductance and activation kinetics. Likewise, the off-path closed state was observed in simulations, but its physiological relevance remains to be established experimentally. Future structural and functional studies, including targeted mutagenesis of the lipid-interacting residues and characterization of lipid-dependent receptor activity, could further test the proposed coupling mechanism.

More broadly, the gating features resolved here provide a structural framework for exploring chemical modulation of mosquito odorant receptors. The orthosteric pocket, the M288-to-S6 connection, and the S6-S7b interface represent distinct regions through which ligand binding could influence receptor conformation and function. These sites could therefore be explored in structure-guided searches for antagonists or allosteric modulators that stabilize nonconducting or otherwise altered conformational states. For AaegOR10 and related mosquito receptors, such compounds could subsequently be tested for effects on odorant receptor activity and, ultimately, odor-guided behaviors relevant to host seeking.

## Methods

### Modeling of the apo and ligand-bound states of AaegOR10

The cryo-EM structures of AaegOR10 in its apo state (AaegOR10_Apo-Closed_), and its complexes with o-cresol (AaegOR10_Holo-Open_) (PDB IDs: 8V00 and 8V02),^15^ were used as the templates to build the atomistic models. To systematically decouple the influence of ligand binding from the receptor’s gating conformation, we also constructed two additional systems: AaegOR10_Apo-Open_ and AaegOR10_Holo-Closed_. The AaegOR10_Apo-Open_ system was generated by the in silico removal of *o*-cresol from the ligand-bound open-state cryo-EM structure. Conversely, the AaegOR10_Holo-Closed_ system was modeled by incorporating the ligand into the binding pocket of the apo-closed structure, utilizing the ligand’s coordinates from the experimental bound structure as a spatial reference for alignment.

Missing loop regions were modeled using the SWISS-MODEL web server,^35^ and the N- and C-termini of the full-length structure were patched with standard terminus patches (NTER or CTER) using the standard CHARMM-GUI procedure. The protonation states of all titratable residues were determined at pH 7.0 using PROPKA 3.0 via the CHARMM-GUI platform.^36^ All residues were maintained in their standard protonation states, and histidine residues were modeled in the tautomeric form with protonation at the *N_δ_* position. These models were then embedded in a 1-palmitoyl-2-oleoyl-sn-glycero-3-phosphocholine (POPC) bilayer and solvated in a cubic water box containing 0.15 M NaCl. The water box dimensions were 12.0 nm × 12.0 nm × 14.0 nm, resulting in approximately 210,000 atoms per system. Membrane insertion of AaegOR10 was predicted using OPM (Orientations of Proteins in Membranes) version 2.0.^37^ System construction was carried out using the CHARMM-GUI web server.^38^ The CHARMM36m force field was applied to the proteins,^39^ and the TIP3P model was used for water.^40^ The CHARMM-WYF parameters were utilized for cation-*π* interactions, and the ligand was parameterized using the CGenFF force field.^41^

In the MD simulations, the temperature was maintained at 310 K using a V-rescale thermostat with a 1 ps coupling constant, and the pressure was maintained at 1.0 bar using a Parrinello-Rahman barostat with a 5 ps coupling constant. A cutoff of 1.2 nm was applied to the van der Waals interactions using a switching function starting at 1.0 nm. The cutoff for short-range electrostatic interactions was also set to 1.2 nm, and long-range electrostatic interactions were calculated using the particle mesh Ewald algorithm with a mesh spacing of 0.12 nm. A reciprocal grid of 96 × 96 × 129 cells was used with fourth-order B-spline interpolation. All simulations were performed using a GPU-accelerated version of GROMACS 2025.^42^

### Unbiased MD simulation and analysis

Following initial energy minimization using the steepest descent algorithm, a six-step equilibration process was performed with gradually released positional restraints. Production MD simulations were then conducted under semi-isotropic NPT conditions. Hydrogen mass repartitioning^43^ enabled a 2.5 fs time step, and each simulation was run for 1000 ns. Three independent 1000 ns unbiased MD replicas were performed for each starting conformation, giving 6 *µ*s for the apo ensemble and 6 *µ*s for the holo ensemble.

These 3 *µ*s of unbiased trajectories for each system served as the basis for initial conformational and interaction analyses. Specifically, the probability distributions of the collective variables (CVs) were calculated using PLUMED 2.9^44^ to evaluate the sampling breadth of the initial ensembles. Long-range allosteric communication and signaling pathways were explored using MDPath,^28^ while the dynamic residue-residue interaction networks, including hydrophobic interactions, hydrogen bonds and van der Waals contacts, were quantified using the GetContacts software.^45^

### Co-evolutionary Analysis

To investigate the evolutionary constraints maintaining the mechanical coupling between transmembrane helices, we performed a co-evolutionary analysis using the EVcouplings framework (https://evcouplings.org/).^46^ The amino acid sequence of the OR chain of AaegOR10 was used as the query to identify conserved residue-residue interactions. Multiple sequence alignments (MSAs) were generated, and evolutionary couplings were calculated based on direct coupling analysis (DCA).^47^ For the identification of functionally significant residue pairs, we applied a confidence cutoff of 0.3.

### Feature space selection

To construct an informative and computationally efficient MSM, we implemented a hierarchical feature selection protocol. The initial feature pool was designed to monitor the global gating motions and local rearrangements of the S5-S7 module. The final set of features was distilled through both structural geometric filtering and trajectory-based statistical analysis.

1. Inter-subunit interactions (34 features): To capture the collective opening of the heteromeric pore, we initially calculated 24 minimum pairwise distances between the OR *S*5-*S*7 helices and the geometric center of the *S*1-*S*6 bundle of each Orco subunit for each of the three OR-Orco interfaces, yielding 72 candidate distances. These features were then filtered based on the Jensen-Shannon divergence (JSD) of the unbiased MD distributions, and 34 distance features with JSD *>* 0.2 were retained as descriptors of inter-subunit gating motions.^48^
2. Intra-subunit Coordination (268 features): We initially considered all residue pairwise distances between the *S*5-*S*7 module and the *S*1-*S*4 bundle within the OR subunit. A two-tier filtering strategy was applied: first, we performed a structure-based screening using the apo and bound cryo-EM models, retaining only those pairs with a minimum distance change *>* 10 ^°^A between the two states and where at least one state exhibited a distance *<* 30 ^°^A. Subsequently, we employed these unbiased MD simulations and used the same trajectory-based JSD analysis to further validate the candidate distances.
3. Torsional Dynamics (118 features): We incorporated 118 dihedral angles, comprising the side-chain dihedrals (*χ*) of the *S*5-*S*7 helices and backbone dihedrals within the flexible loop regions. These features were filtered using the same trajectory-based JSD analysis to prioritize residues involved in local conformational switches during activation.

In total, 420 features (34 OR-Orco distances, 268 OR-OR distances, and 118 dihedrals) were selected by PLUMED 2.9^44^ to define the high-dimensional conformational space for dimensionality reduction and MSM construction (Fig. S13).

### Adaptive Sampling

To efficiently explore the high-dimensional transition landscape and bridge the millisecond-scale gating kinetics, we implemented an extensive adaptive sampling protocol. For the ligand-bound system (AaegOR10_Holo_), sampling was initiated from 30 structures generated using the PyMOL morphing function, each subjected to 1 *µ*s of MD.^49^ We then performed 20 rounds of adaptive sampling. Features were extracted every nanosecond from both the initial morphing round and subsequent adaptive sampling rounds. At each round, the accumulated holo trajectories were used to update the feature-based tICA transformation.^50,51^ The resulting tIC projections were clustered into 200 microstates using the kNN (k-nearest neighbors)-dPCA algorithm,^52^ and an MSM was estimated for that round. New seeds were selected using a macrocount-based discovery metric to prioritize undersampled regions of phase space.^53,54^ Each round consisted of 12 parallel trajectories, yielding approximately 270 *µ*s in total.

For the apo system (AaegOR10*_apo_*), we adopted a transfer-learning-inspired strategy to accelerate convergence. The projected holo conformations were partitioned into 50 microstates using kNN-dPCA, with the resulting cluster centers serving as initial seeds to promote robust MSM connectivity while limiting the computational cost. Following a 0.25 *µ*s equilibration phase for each seed, two independent 1-*µ*s simulations were initiated. During subsequent adaptive sampling, the accumulated apo trajectories were projected onto the common holo-derived tICA coordinate space, and new seeds were iteratively selected using the same macrocount-based discovery metric. Three additional rounds of adaptive sampling were performed, each comprising 12 parallel 1-*µ*s trajectories, bringing the cumulative sampling time to 136 *µ*s.

Additionally, we incorporated six previously generated 1 *µ*s trajectories into each ensemble, comprising three replicas from each of the open and closed starting conformations. The final datasets therefore comprised 276 *µ*s of aggregate simulation time for the holo system and 142 *µ*s for the apo system (Table S7).

### Construction of Markov State Models (MSMs) and Bayesian MSMs (BMSMs)

Markov state models were constructed separately for the holo and apo ensembles to quantify the thermodynamics and kinetics of the gating process. ^26,27^ The 420-dimensional feature space was reduced using time-lagged independent component analysis (tICA) with a 20 ns lag time.^50,51^ For the final MSM analysis, the tICA transformation was trained on the complete holo dataset and then applied to both the holo and apo trajectories, placing the two ensembles in a common kinetic coordinate space.

- During the adaptive sampling phase, we retained a sufficient number of tICs to account for 95% of the total kinetic variance, ensuring a broad exploration of the conformational space.
- For the final MSM construction, we specifically utilized the first 8 tICs. This deliberate selection was aimed at enhancing the connectivity between metastable states and improving the robustness of the kinetic transition network, as these top components capture the slowest and most biologically relevant allosteric movements.

Microstates were then defined independently for the holo and apo ensembles by clustering their respective projections in the common holo-derived tICA space using kNN-dPCA. The general concept of the clustering approach is consistent with that described in the work of Liu et al.;^52^ however, to enhance flexibility, a custom-written script was employed in the present study. The nearest-neighbor count *k* in kNN was set to 10,000, and the distance cutoff *d_c_* in dPCA was set to 2. Each ensemble was clustered separately into 200 microstates to construct its corresponding MSM. The MSMs were estimated at a lag time (*τ* ) of 200 ns, as supported by convergence of the implied timescales.

To evaluate the statistical uncertainty of the MSMs, we employed Bayesian MSMs (BMSMs) to generate an ensemble of 100 transition-matrix samples for each ensemble.^55^ These samples were used to estimate confidence intervals for the thermodynamic and kinetic observables. Specifically, the reported mean values, including free-energy estimates and transition timescales, were derived from the maximum-likelihood MSMs. The associated statistical errors were estimated as the standard errors across the 100 transition-matrix samples in each BMSM ensemble. Model robustness was assessed using the Chapman-Kolmogorov (CK) test.^27^ The deeptime package was used for tICA, MSM/BMSM construction, and subsequent analyses.^56^

### Thermodynamic and Kinetic Characterization of the Gating Landscape based on MSM/BMSM

To resolve the properties of the transition processes, we employed a combination of thermodynamic and kinetic analysis based on the constructed MSM/BMSM.

The FES was projected onto the first two tIC dimensions using MSM-based Koopman reweighting.^57^ We then computed the minimum free-energy path (MFEP) from the two-dimensional tIC FES and projected the FES onto this pathway to obtain a corresponding one-dimensional free-energy profile using custom scripts. To resolve metastable states, we employed the Perron cluster cluster analysis (PCCA+) algorithm^58^ to discretize the microstate space into macrostates. The apo ensemble was resolved into five macrostates, whereas the ligand-bound ensemble was resolved into six, including the additional off-path intermediate.

We calculated the mean first-passage times (MFPTs) between the resolved macrostates to characterize the global kinetics of the gating process. ^59,60^ By constructing an MFPT-based kinetic network, we identified the rate-limiting steps and the kinetic signatures governing the receptor’s transition landscape.

Furthermore, we performed two complementary TPT analyses.^26,48^ First, separate pairwise TPT calculations were performed for each ordered pair of macrostates. Then, to characterize the dominant gating routes in both directions, pathway decompositions were performed from each open-like source macrostate to C and from C or C*^′^* to O. For each source-target pair, the total reactive flux was decomposed into individual pathways, which were ranked according to their pathway flux. The pathway fraction was calculated as the pathway flux divided by the total reactive flux for the corresponding source-target pair. All MFPTs, fluxes and pathway fractions were calculated using the maximum-likelihood MSMs, with SE estimated across the 100 transition-matrix samples in the corresponding BMSM ensemble.

### Comparative sequence and structural analysis of *Aedes* odorant receptors

To complement the MD-based analysis, we examined the sequence and structural conservation of the kink-prone region across odorant receptor systems from the genus *Aedes*. Odorant receptor protein sequences were retrieved from VectorBase.^29^ Only sequences from the genus *Aedes* were retained, including proteins longer than 300 amino acids and shorter than 450 amino acids with an unambiguous UniProt identifier. OR and Orco sequences were classified separately.

For each retained OR sequence, an AlphaFold3 input was generated to model an OR-Orco tetramer.^30^ Each tetramer comprised one OR subunit and three identical Orco subunits. OR subunits containing six to eight transmembrane helices were retained, while structures that were clearly inconsistent with the expected OR architecture were removed. The resulting complexes were aligned using FoldMason.^31^ The alignment was mapped to the residue numbering of AaegOR10, and the region corresponding to S291-H299 was extracted for comparative analysis. For each aligned position, the corresponding Chou-Fasman *α*-helix propensity value was assigned and summarized across the retained *Aedes* systems.^61^

### Functional characterization of AaegOR10 and its mutants

The full-length sequence of AaegOR10 was synthesized by GENEWIZ, Inc. (Suzhou, China), and AaegOrco was amplified from *A. aegypti* using gene-specific primers. Both coding sequences were subsequently cloned into the pT7TS vector. Site-directed mutagenesis was performed by PCR on the entire clone using mutant primers and Phusion High-Fidelity DNA Polymerase (NEB, Ipswich, MA, USA). The primer sequences were listed in Table S8.

The protocols of cRNA synthesis, *Xenopus* oocytes preparation, cRNA injection and two-electrode voltage-clamp (TEVC) recording were previously described.^62^ The cRNAs were synthesized using the mMESSAGE mMACHINE T7 kit (Invitrogen, Austin, TX, USA) according to the manufacturer’s instructions and resuspended in RNase-free water at 2 *µ*g/*µ*L. The cRNAs of AaegOR10 or each of its mutants were co-injected with AaegOrco into mature healthy *Xenopus* oocytes (27.6 nL per oocyte). The injected oocytes were incubated in 1 *×* Ringer’s buffer at 18 *^◦^*C for 3-6 days.

TEVC recording was performed to measure the whole-cell currents elicited by *o*-cresol, a well-established ligand of AaegOR10.^62^ *o*-Cresol (CAS No. 95-48-7; *>*99% purity) was purchased from TCI (Tokyo, Japan), and a 1 M stock solution was prepared in DMSO and stored at *−*20*^◦^*C. Working solutions at the desired concentrations were freshly prepared in 1*×* Ringer’s buffer before each experiment. Oocytes co-expressing wild-type or mutant AaegOR10 with AaegOrco were stimulated with *o*-cresol at concentrations ranging from 10*^−^*^8^, 10*^−^*^7^, 10*^−^*^6^, 10*^−^*^5^, 10*^−^*^4^, and 10*^−^*^3^ M. The signals were recorded using an OC-725D amplifier (Warner Instruments, Hamden, CT, USA) at a holding potential of *−*80 mV. Data acquisition and analysis were performed using Axon Digidata 1550B and pCLAMP10.6 software (Molecular Devices, San Jose, CA, USA). Dose-response curves were fitted using a custom-written script with a three-parameter Hill equation.

### Data visualization and representation

Trajectory and structure visualization was conducted using PyMOL.^49^ Plots were generated using Matplotlib 3.7 and Python 3.10.

## Data availability

Analysis scripts are available on GitHub at https://github.com/TiefengSong/AaegOR10. Simulation inputs, MD trajectories, predicted *Aedes* OR-Orco structural models and electrophysiological data are available at https://ngdc.cncb.ac.cn/omix/release/OMIX021586.

## Supporting Information

The Supporting Information is available free of charge at [Pending upon publication].

Simulation systems and analytical scope, state populations and transition pathways, sampling and primer information, electrophysiological data, and additional interaction, kinetic, and structural analyses.

## Acknowledgement

Y.W. acknowledges the financial support of the Zhejiang Provincial National Science Foundation of China (No. LZ24C050003), the National Natural Science Foundation of China (No. 32371300), computational support of the Information Technology Center and State Key Lab of CAD&CG at Zhejiang University, and the Postdoctoral Fellowship Program of CPSF (No. GZC20252512). ChatGPT was used to improve the grammar, consistency, and readability of the manuscript.

