## Supplementary material for "From Ligand Binding to Pore Opening in the Mosquito Odorant Receptor AaegOR10": SI

### Supporting Information for: From Ligand Binding to Pore Opening in the Mosquito Odorant Receptor AaegOR10

### Supplementary Information

**Table S1:** Summary of biological insights obtained from different simulation systems.

| System | Binding Mode | Interaction Mode | Free Energy Surface | Metastable State | Transition Pathway and Kinetics |
|---|---|---|---|---|---|
| AaegOR10$_{\text{Holo-Open}}$ | Yes | Yes | Yes | Yes | Yes |
| AaegOR10$_{\text{Apo-Closed}}$ | - | Yes | Yes | Yes | Yes |

**Table S2:** Stationary populations of the macrostates in the holo and apo ensembles. Populations were obtained from the corresponding maximum-likelihood MSM.

| Ensemble | Macrostate | Population (%) |
|---|---|---|
| Holo | O | 18.83 |
| | O′ | 0.72 |
| | I | 21.50 |
| | I′ | 8.66 |
| | C | 26.77 |
| | C′ | 23.52 |
| Apo | O | 10.53 |
| | O′ | 4.43 |
| | I | 5.81 |
| | C | 41.18 |
| | C′ | 38.05 |

**Table S3:** Selected TPT pathways to the closed state in the holo ensemble. Pathway fractions denote the fraction of the total source-to-C reactive flux carried by each pathway. Values are from the maximum-likelihood MSM, with SE estimated across 100 BMSM transition-matrix samples.

| Start | Target | Pathway | Pathway reactive flux ($ms^{-1}$) | Pathway fraction (%) |
|---|---|---|---|---|
| O′ | C | O′ → I → C | $0.93 \pm 0.03$ | $65.45 \pm 0.32$ |
| | | O′ → I → O → C | $0.33 \pm 0.01$ | $23.24 \pm 0.50$ |
| | | O′ → I → I′ → C | $0.09 \pm 0.00$ | $6.40 \pm 0.08$ |
| | | O′ → I → C′ → C | $0.03 \pm 0.00$ | $2.20 \pm 0.08$ |
| | | O′ → O → C | $0.02 \pm 0.01$ | $1.30 \pm 0.42$ |
| I′ | C | I′ → I → C | $9.12 \pm 0.09$ | $57.50 \pm 0.32$ |
| | | I′ → C | $3.49 \pm 0.05$ | $22.02 \pm 0.28$ |
| | | I′ → I → O → C | $2.07 \pm 0.04$ | $13.07 \pm 0.26$ |
| | | I′ → O → C | $0.82 \pm 0.02$ | $5.19 \pm 0.14$ |
| | | I′ → I → C′ → C | $0.25 \pm 0.01$ | $1.58 \pm 0.06$ |
| O | C | O → I → C | $11.86 \pm 0.13$ | $47.99 \pm 0.42$ |
| | | O → C | $10.58 \pm 0.21$ | $42.80 \pm 0.48$ |
| | | O → I′ → C | $0.90 \pm 0.01$ | $3.62 \pm 0.06$ |
| | | O → I → I′ → C | $0.54 \pm 0.01$ | $2.18 \pm 0.06$ |
| | | O → C′ → C | $0.42 \pm 0.03$ | $1.70 \pm 0.15$ |

**Table S4:** Selected TPT pathways to the closed state in the apo ensemble. Pathway fractions denote the fraction of the total source-to-C reactive flux carried by each pathway. Values are from the maximum-likelihood MSM, with SE estimated across 100 BMSM transition-matrix samples.

| Start | Target | Pathway | Pathway reactive flux ($ms^{-1}$) | Pathway fraction (%) |
|---|---|---|---|---|
| O′ | C | O′ → I → C | $1.09 \pm 0.04$ | $58.22 \pm 0.46$ |
| | | O′ → O → I → C | $0.66 \pm 0.03$ | $35.14 \pm 0.80$ |
| | | O′ → O → I → C′ → C | $0.10 \pm 0.01$ | $5.32 \pm 0.38$ |
| | | O′ → O → C | $0.03 \pm 0.02$ | $1.32 \pm 0.72$ |
| O | C | O → I → C | $1.55 \pm 0.05$ | $76.53 \pm 0.83$ |
| | | O → O′ → I → C | $0.34 \pm 0.02$ | $16.71 \pm 0.55$ |
| | | O → O′ → I → C′ → C | $0.11 \pm 0.01$ | $5.34 \pm 0.38$ |
| | | O → C | $0.03 \pm 0.03$ | $1.43 \pm 0.92$ |

**Table S5:** Selected TPT pathways to the open state in the holo ensemble. Pathway fractions denote the fraction of the total source-to-O reactive flux carried by each pathway. Values are from the maximum-likelihood MSM, with SE estimated across 100 BMSM transition-matrix samples.

| Start | Target | Pathway | Pathway reactive flux ($ms^{-1}$) | Pathway fraction (%) |
|---|---|---|---|---|
| C | O | $C \to I \to O$ | $11.86 \pm 0.13$ | $47.99 \pm 0.42$ |
| | | $C \to O$ | $10.58 \pm 0.21$ | $42.80 \pm 0.48$ |
| | | $C \to I' \to O$ | $0.90 \pm 0.01$ | $3.62 \pm 0.06$ |
| | | $C \to I' \to I \to O$ | $0.54 \pm 0.01$ | $2.18 \pm 0.06$ |
| | | $C \to C' \to O$ | $0.42 \pm 0.03$ | $1.70 \pm 0.15$ |
| C′ | O | $C' \to C \to I \to O$ | $7.41 \pm 0.07$ | $46.19 \pm 0.45$ |
| | | $C' \to C \to O$ | $7.10 \pm 0.13$ | $44.26 \pm 0.52$ |
| | | $C' \to I \to O$ | $0.51 \pm 0.02$ | $3.17 \pm 0.10$ |
| | | $C' \to C \to I' \to O$ | $0.48 \pm 0.01$ | $2.98 \pm 0.05$ |
| | | $C' \to O$ | $0.43 \pm 0.03$ | $2.66 \pm 0.22$ |

**Table S6:** Selected TPT pathways to the open state in the apo ensemble. Pathway fractions denote the fraction of the total source-to-O reactive flux carried by each pathway. Values are from the maximum-likelihood MSM, with SE estimated across 100 BMSM transition-matrix samples.

| Start | Target | Pathway | Pathway reactive flux ($ms^{-1}$) | Pathway fraction (%) |
|---|---|---|---|---|
| C | O | $C \to I \to O$ | $1.55 \pm 0.05$ | $76.53 \pm 0.83$ |
| | | $C \to I \to O' \to O$ | $0.34 \pm 0.02$ | $16.71 \pm 0.55$ |
| | | $C \to C' \to I \to O' \to O$ | $0.11 \pm 0.01$ | $5.34 \pm 0.38$ |
| | | $C \to O$ | $0.03 \pm 0.03$ | $1.43 \pm 0.92$ |
| C′ | O | $C' \to C \to I \to O$ | $1.12 \pm 0.03$ | $76.25 \pm 0.97$ |
| | | $C' \to C \to I \to O' \to O$ | $0.19 \pm 0.01$ | $12.74 \pm 0.77$ |
| | | $C' \to I \to O' \to O$ | $0.13 \pm 0.01$ | $9.13 \pm 0.67$ |
| | | $C' \to C \to O$ | $0.03 \pm 0.03$ | $1.88 \pm 1.17$ |

**Table S7:** Summary of aggregate simulation sampling and its composition.

| **Ensemble** | **Sampling mode** | **Aggregate sampling ($\mu$s)** | **Composition** |
|---|---|---|---|
| AaegOR10$_{\text{Holo}}$ | Unbiased MD | 6 | 3 × 1 $\mu$s from AaegOR10$_{\text{Holo-Open}}$ and 3 × 1 $\mu$s from AaegOR10$_{\text{Holo-Closed}}$ |
| AaegOR10$_{\text{Holo}}$ | Adaptive sampling | 270 | 30 initial 1-$\mu$s trajectories followed by 20 rounds of 12 × 1-$\mu$s trajectories |
| AaegOR10$_{\text{Apo}}$ | Unbiased MD | 6 | 3 × 1 $\mu$s from AaegOR10$_{\text{Apo-Open}}$ and 3 × 1 $\mu$s from AaegOR10$_{\text{Apo-Closed}}$ |
| AaegOR10$_{\text{Apo}}$ | Adaptive sampling | 136 | 100 initial 1-$\mu$s trajectories followed by 3 rounds of 12 × 1-$\mu$s trajectories |
| AaegOR10$_{\text{Holo}}$ | Total | 276 | 270 $\mu$s adaptive sampling plus 6 $\mu$s unbiased MD |
| AaegOR10$_{\text{Apo}}$ | Total | 142 | 136 $\mu$s adaptive sampling plus 6 $\mu$s unbiased MD |

**Table S8:** Primer sequences used for cloning and site-directed mutagenesis.

| Primer name | Purpose | Sequence (5'-3') |
|---|---|---|
| AaegOR10-F | Cloning/amplification | ATGGCAAGCATTCTTGATTGC |
| AaegOR10-R | Cloning/amplification | TTAATTATAAACCCGACGCAG |
| AaegOR10-M288A-F | Site-directed mutagenesis | CCGCCTACATCTTCGCGATAAT TTCTCA |
| AaegOR10-M288A-R | Site-directed mutagenesis | GCGAAGATGTAGGCGGCGACG |
| AaegOR10-S291G-F | Site-directed mutagenesis | ATCTTCATGATAATTGGTCAAA TCTTCG |
| AaegOR10-S291G-R | Site-directed mutagenesis | CCAATTATCATGAAGATGTAG |
| AaegOR10-I293A-F | Site-directed mutagenesis | ATGATAATTTCTCAAGCCTTCG CCTTCT |
| AaegOR10-I293A-R | Site-directed mutagenesis | GCTTGAGAAATTATCATGAAG |
| AaegOR10-I293Q-F | Site-directed mutagenesis | TGATAATTTCTCAACAATTCGC CTTCT |
| AaegOR10-I293Q-R | Site-directed mutagenesis | TTGTTGAGAAATTATCATGAA |
| AaegOR10-F294A-F | Site-directed mutagenesis | TAATTTCTCAAATCGCCGCCTT CTACTG |
| AaegOR10-F294A-R | Site-directed mutagenesis | GCGATTTGAGAAATTATCATG |
| AaegOR10-F294Q-F | Site-directed mutagenesis | TAATTTCTCAAATCCAAGCCTT CTACTG |
| AaegOR10-F294Q-R | Site-directed mutagenesis | TTGGATTTGAGAAATTATCAT |
| AaegOR10-Y297P-F | Site-directed mutagenesis | CAAATCTTCGCCTTCCCCTGGC ACGCCA |
| AaegOR10-Y297P-R | Site-directed mutagenesis | GGGAAGGCGAAGATTTGAGAA |
| AaegOR10-F356A-F | Site-directed mutagenesis | ATGACGCTGGAAATGGCCCAAT CCCTGC |
| AaegOR10-F356A-R | Site-directed mutagenesis | GCCATTTCCAGCGTCATCGGA |
| AaegOR10-F356Q-F | Site-directed mutagenesis | ATGACGCTGGAAATGCAACAAT CCCTGC |
| AaegOR10-F356Q-R | Site-directed mutagenesis | TTGCATTTCCAGCGTCATCGG |

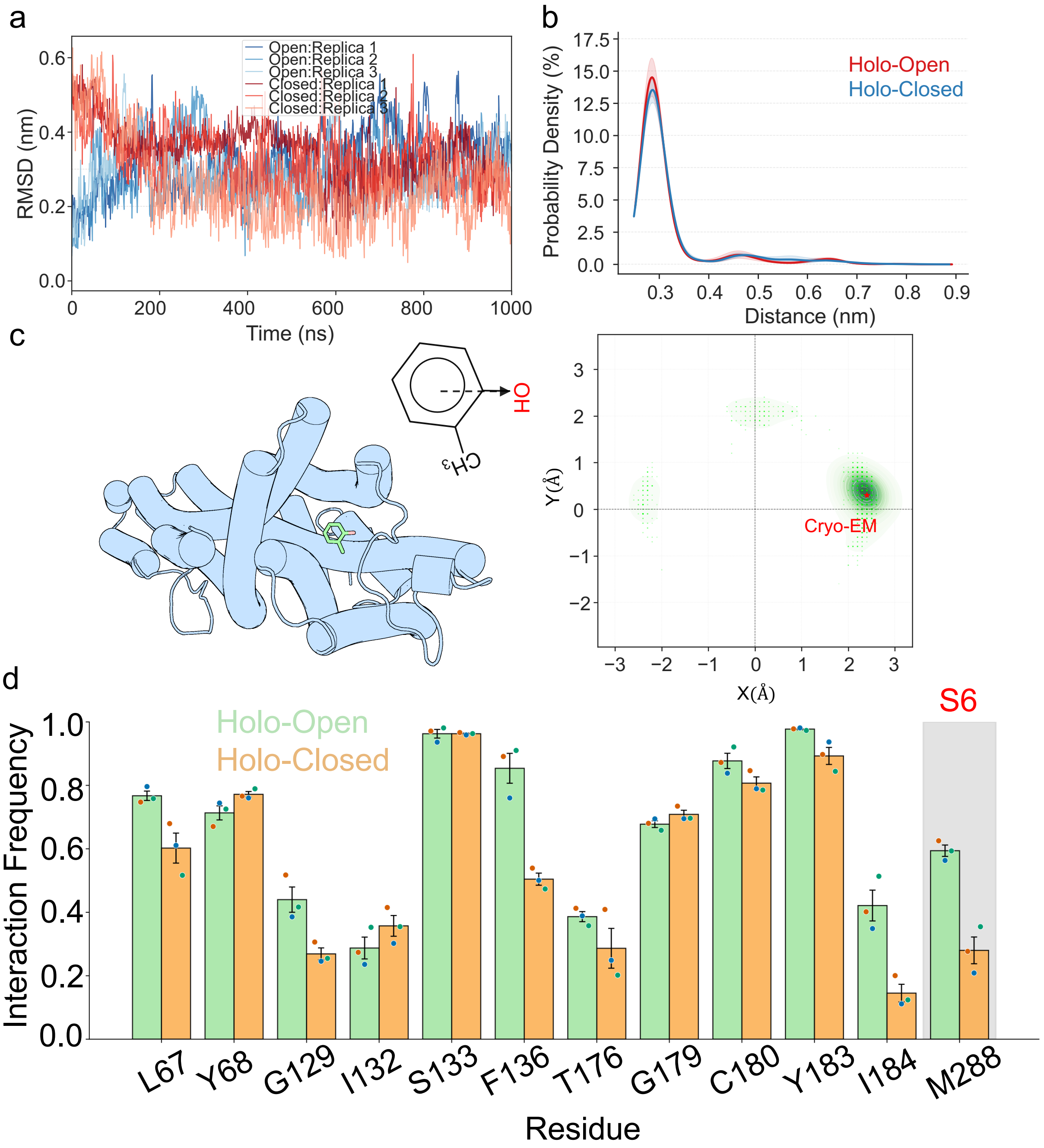


**Figure S1: Ligand RMSD, hydrogen bond interactions and posture distribution.** a) RMSD of o-cresol in unbiased MD of AaegOR10$_{\text{Holo-Open}}$ and **AaegOR10**$_{\text{Holo-Closed}}$. b) The distance between the hydroxyl oxygen of S133 and the hydroxyl oxygen of the ligand in unbiased MD of AaegOR10$_{\text{Holo-Open}}$ and AaegOR10$_{\text{Holo-Closed}}$. Lines show the mean distributions, and shaded regions indicate SE across three trajectories. c) The orientation is defined by the projection of the vector connecting the ligand's para-carbon to its hydroxyl oxygen onto the xy-plane. In its stable binding pose, the ligand is oriented laterally, pointing toward the protein periphery. The left panel provides a structural schematic of the equilibrium ligand posture, while the right panel displays the angular distribution of these orientations across the ensemble. d) Ligand-pocket interaction frequencies. Bars show means, error bars indicate SE, and colored points show individual values from three trajectories.

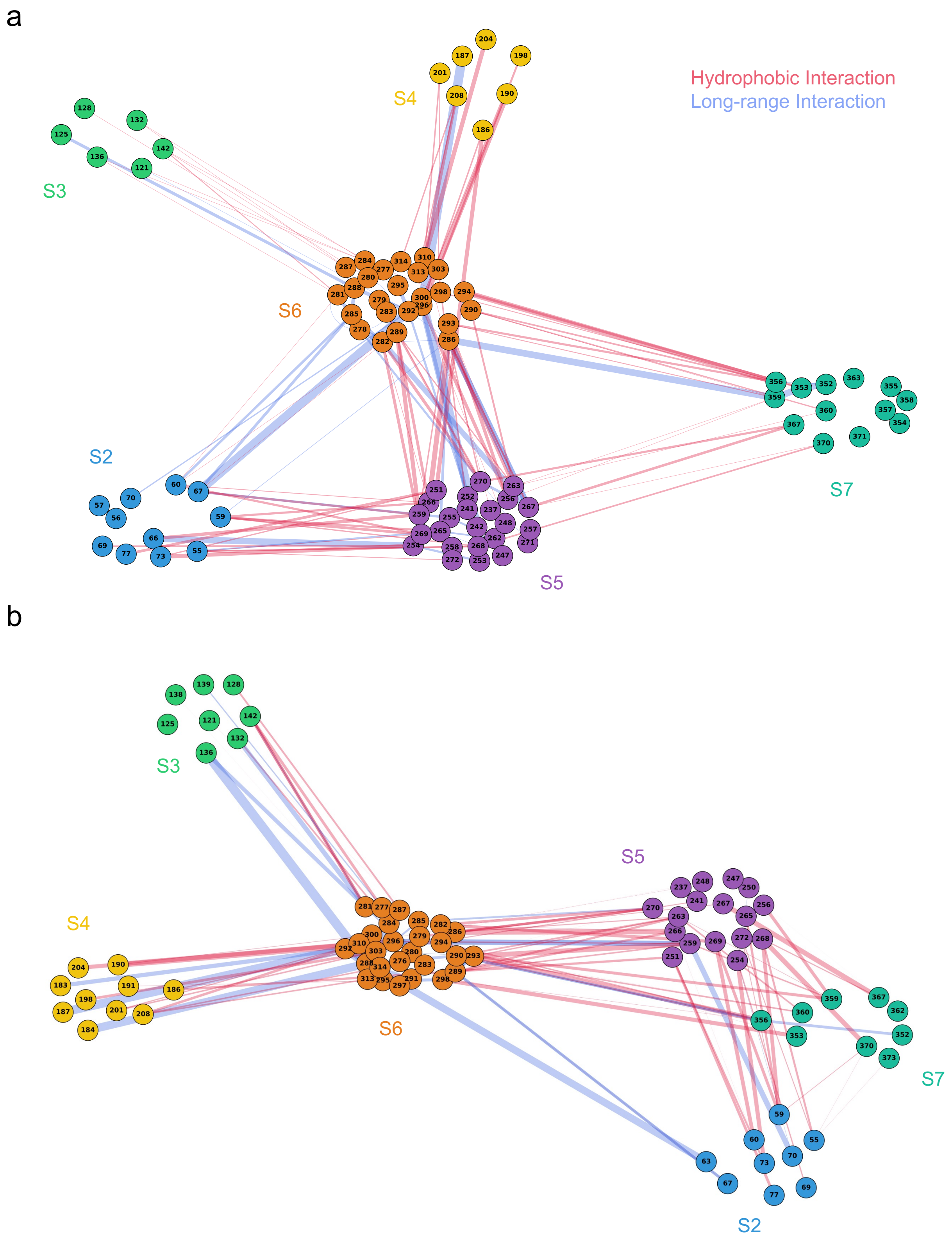


**Figure S2: Interaction networks in the apo-closed and holo-open ensembles.** Long-range (blue lines) and hydrophobic (red lines) interaction networks within AaegOR10$_{\text{Apo-Closed}}$ (a) and AaegOR10$_{\text{Holo-Open}}$ (b), as identified by MDPath. Residues belonging to the same helix are grouped together and represented by consistent colors. The thickness of each line is proportional to the interaction intensity, reflecting the relative strength of the coupling.

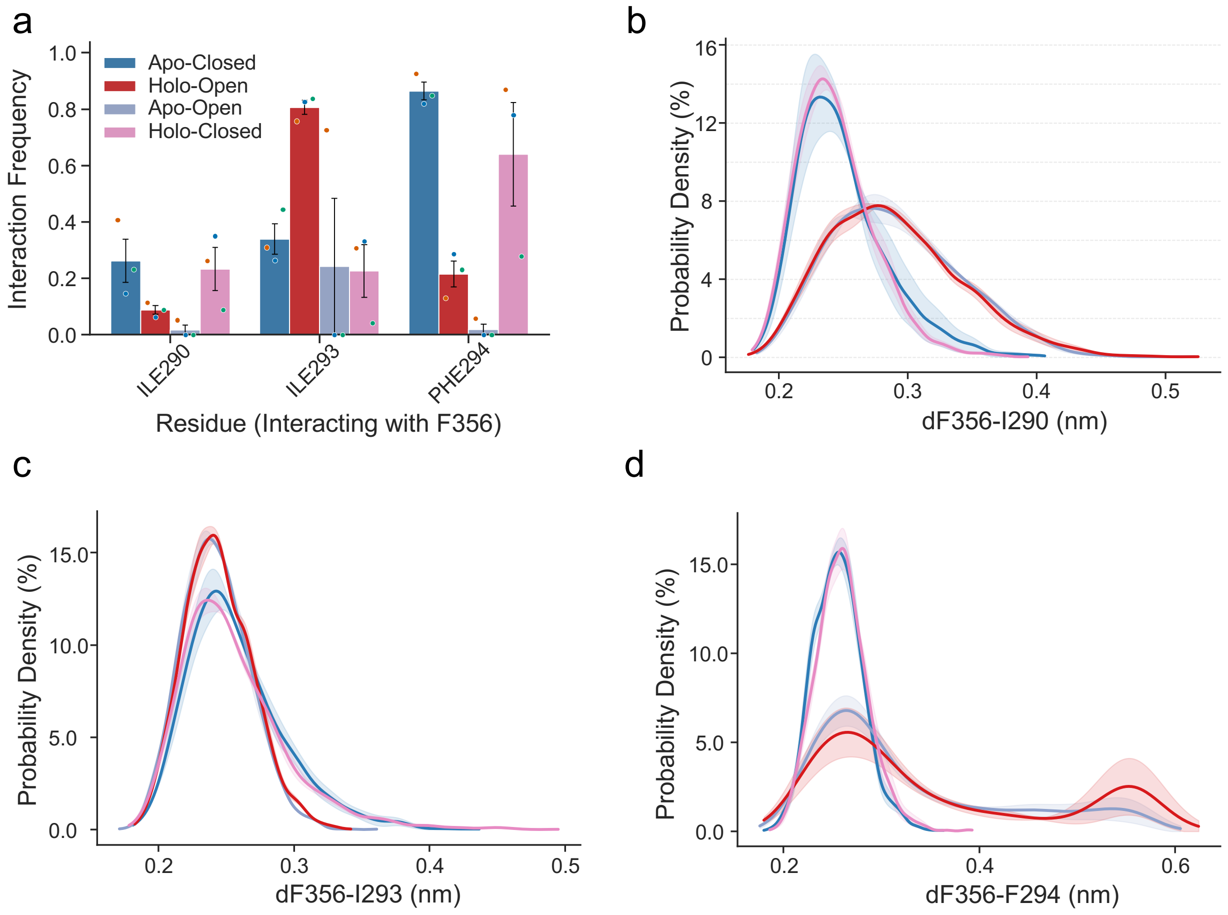


**Figure S3: Hydrophobic interaction analysis at the S6-S7b interface.** a) Hydrophobic interaction frequencies between F356 and I290, I293, and F294 across the four studied systems. Bars show the mean of three trajectories, error bars indicate SE, and colored points show individual trajectory values. b-d) Distance probability distributions between F356 and residues I290 (b), I293 (c), and F294 (d), respectively. Lines show the mean distributions, and shaded regions indicate SE across three trajectories.

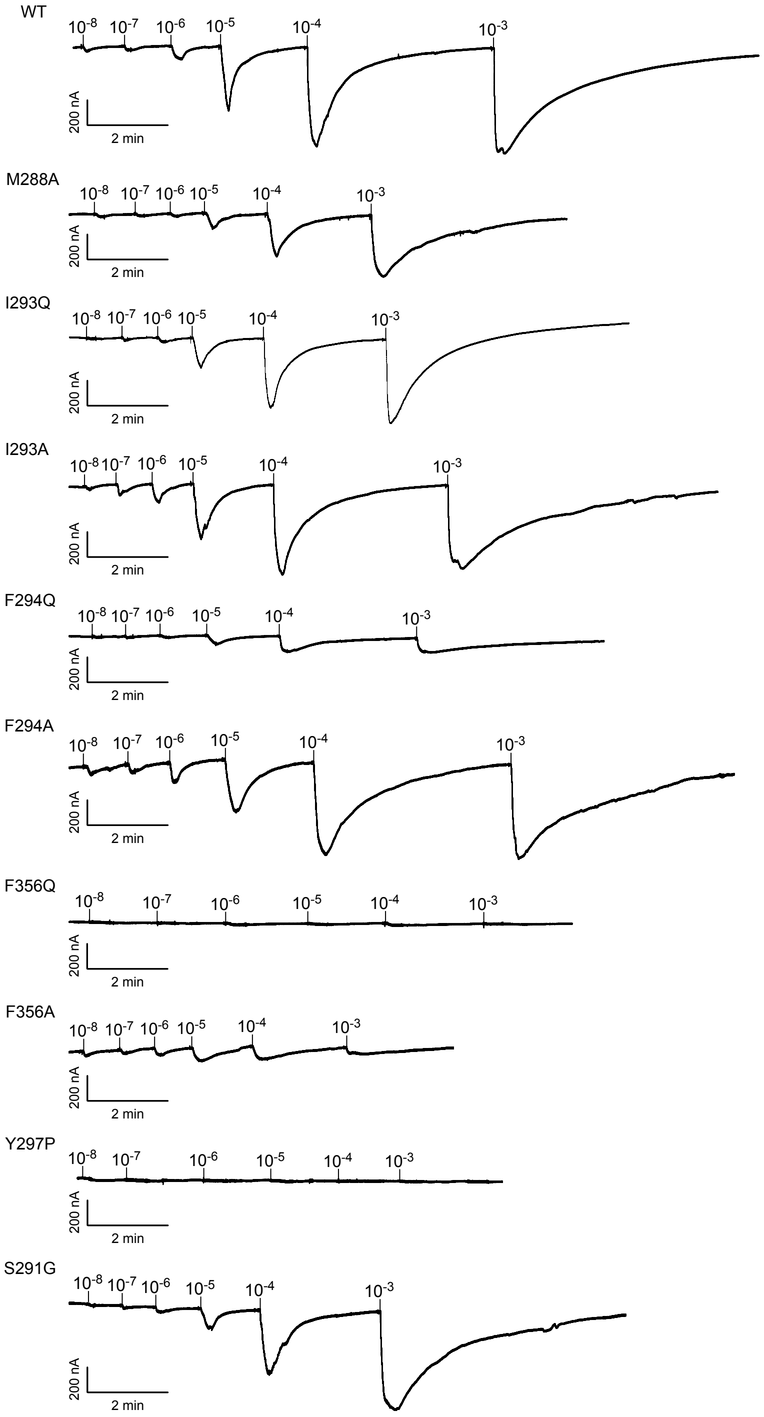


**Figure S4: Representative voltage-clamp current traces of AaegOR10 and its mutants.** Representative current responses recorded during voltage-clamp measurements at increasing *o*-cresol concentrations for wild-type AaegOR10 and the indicated mutants. The traces shown are representative of recordings from each construct; 5 or 6 independent recordings were obtained for each construct. Scale bars indicate 200 nA and 2 min, except where noted.

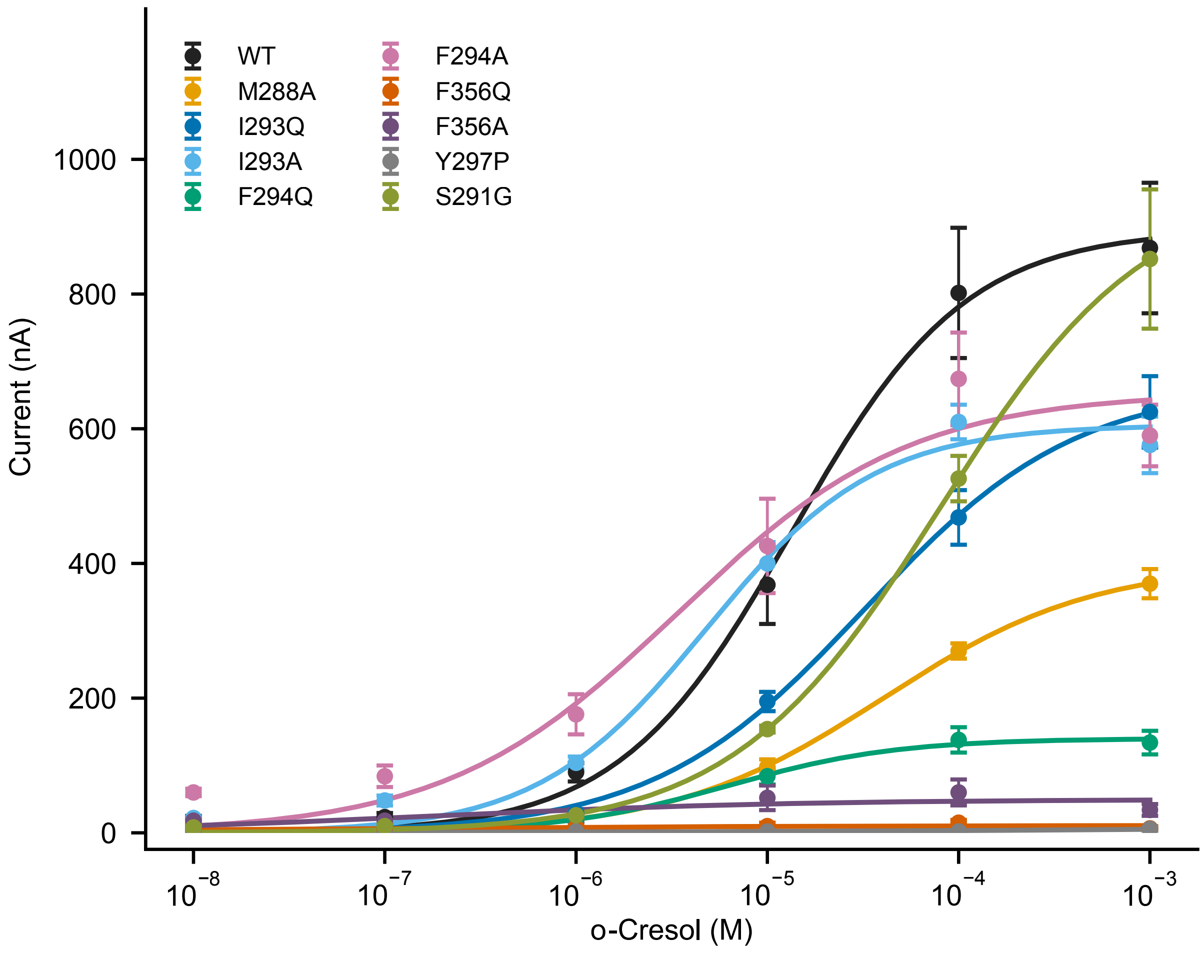


**Figure S5: Raw concentration-response relationships of AaegOR10 and its mutants.** Concentration-dependent current responses measured by voltage clamp for wild-type AaegOR10 and the indicated mutants. Currents are plotted without normalization. Points represent mean values, error bars indicate SE, and each concentration-response relationship was obtained from 5 or 6 independent recordings.

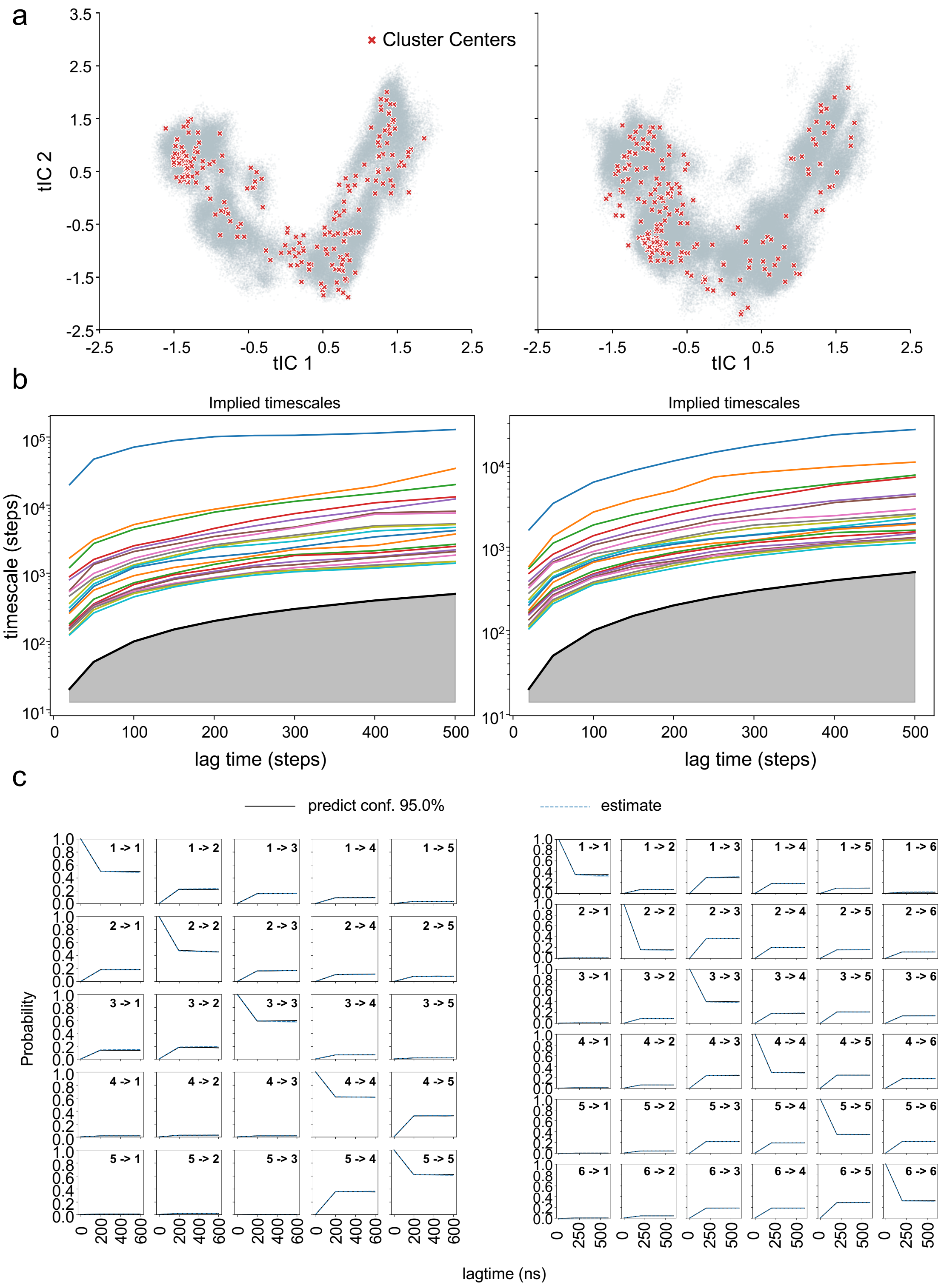


**Figure S6: Validation of the Markov State Models (MSMs).** a) Projection of microstate cluster centers onto the tICA space for the apo (left) and holo (right) ensembles. b) Implied timescales (ITS) as a function of the lag time for the apo (left) and holo (right) MSMs, used to assess model convergence. c) Chapman-Kolmogorov (CK) test results for the apo (left) and holo (right) models, validating the Markovian property of the transitions between macrostates.

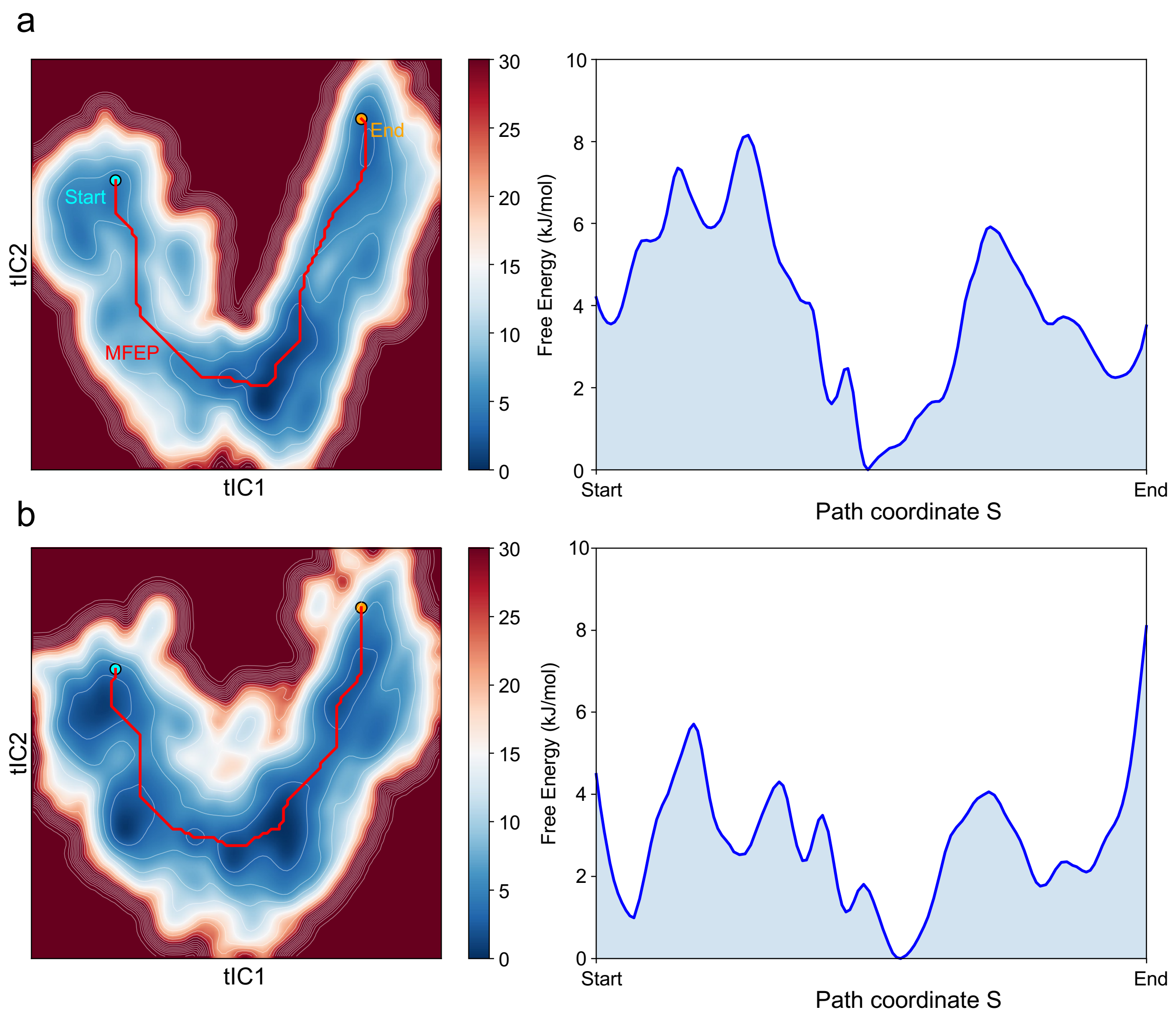


**Figure S7: Minimum free-energy paths and one-dimensional free-energy profiles.** The MFEPs based on the two-dimensional tIC FESs and the corresponding one-dimensional free-energy profiles projected onto these pathways for the apo (a) and holo (b) ensembles.

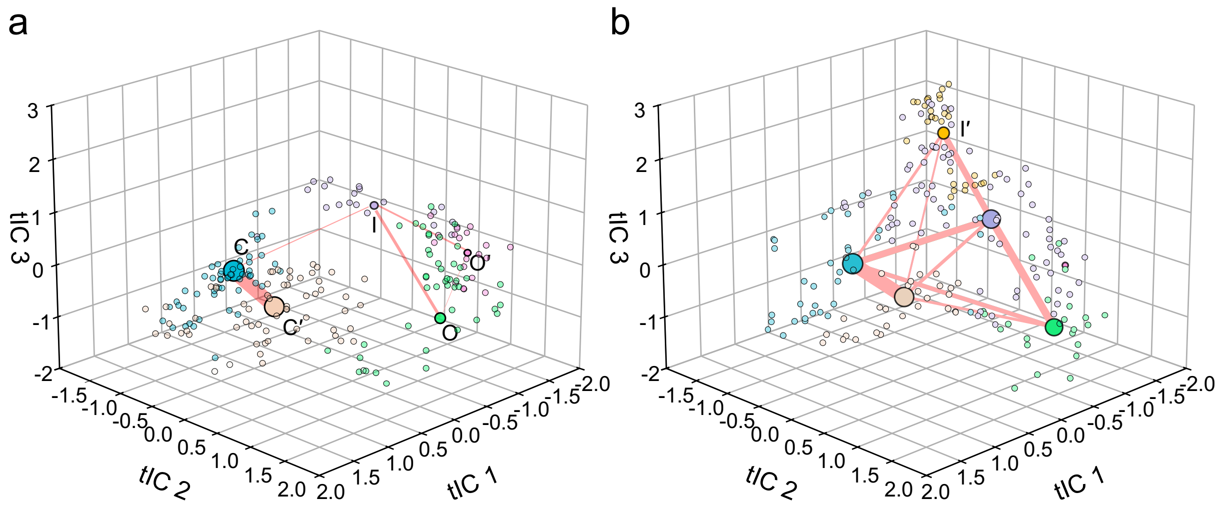


**Figure S8: Microstate distributions and pairwise total reactive fluxes projected onto the first three tICs.** Microstates and coarse-grained macrostates are shown for the apo (a) and holo (b) ensembles. Small points represent microstate cluster centers and are colored according to their macrostate assignments; large labeled circles represent the corresponding macrostates. Separate TPT calculations were performed for each macrostate pair, with line thickness proportional to the corresponding total reactive flux.

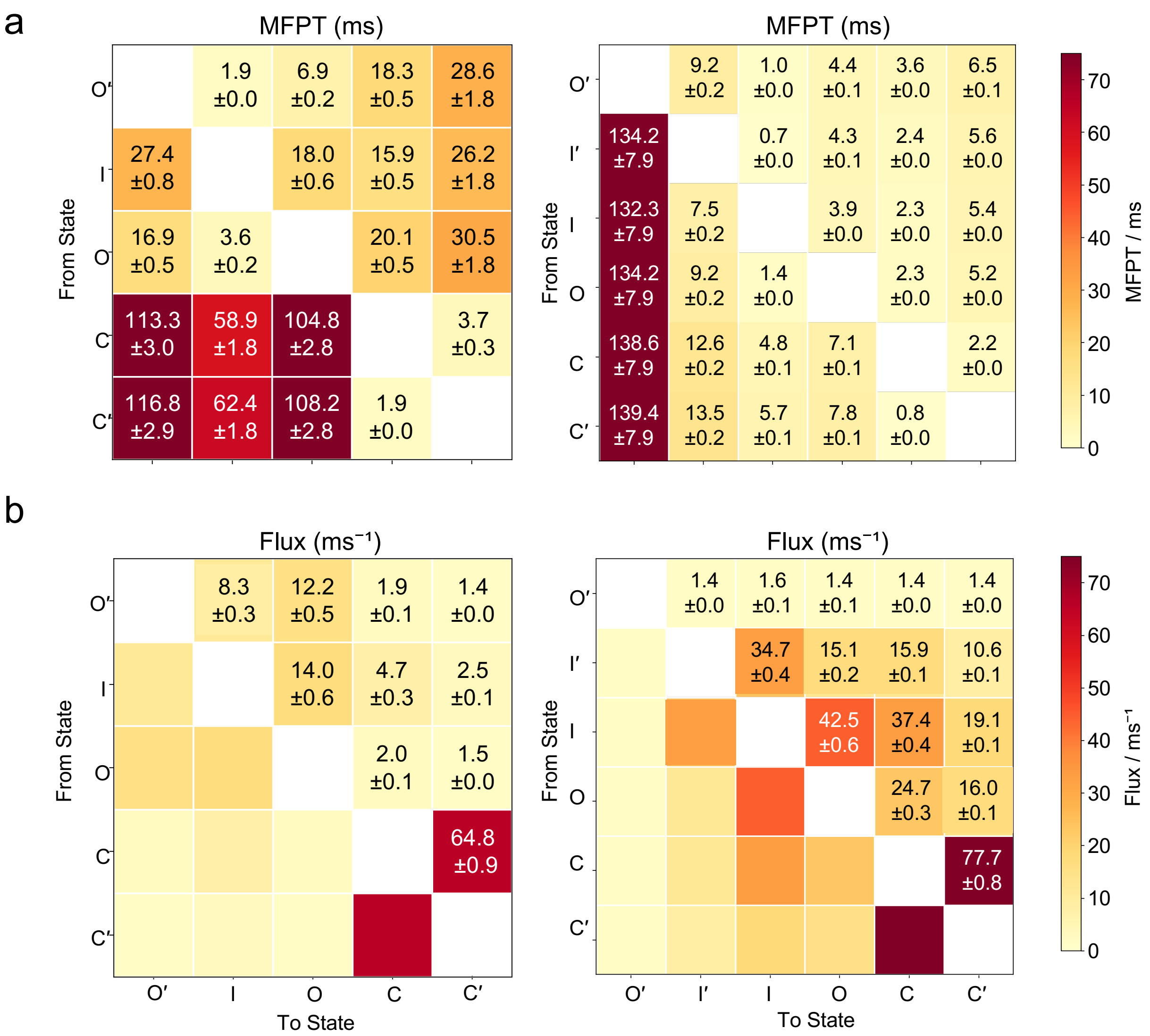


**Figure S9: Kinetic analysis of the macrostate transitions.** a) Mean first-passage times (MFPTs) between the identified macrostates for the apo (left) and holo (right) systems. Numbers indicate MFPTs derived from the maximum-likelihood MSMs, with SE estimated across the 100 transition-matrix samples in the corresponding BMSM ensemble. b) Pairwise total reactive fluxes between the identified macrostates for the apo (left) and holo (right) systems. Values were obtained from separate TPT calculations for each macrostate pair using the maximum-likelihood MSMs, with SE estimated across the 100 transition-matrix samples in the corresponding BMSM ensemble.

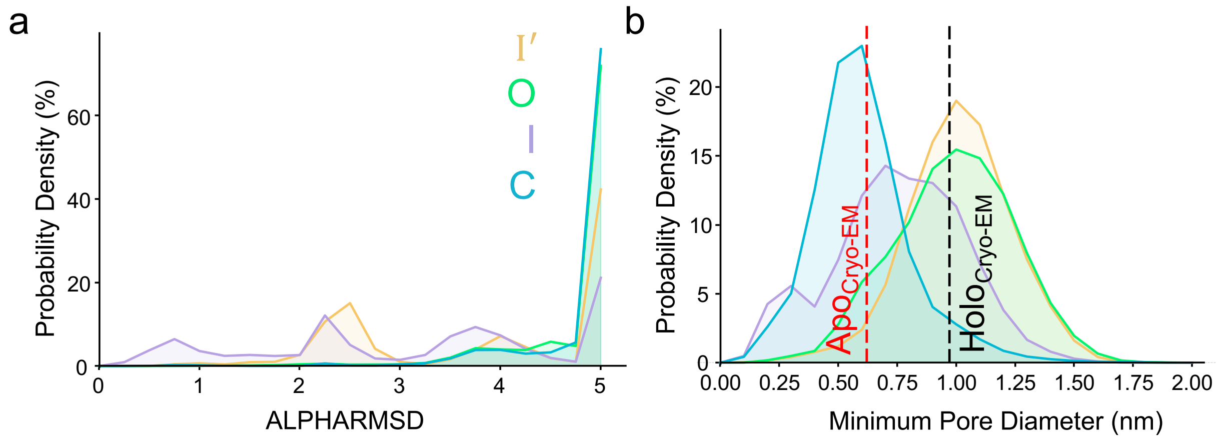


**Figure S10: Structural characteristics of the I and I′ states.** a) Distribution of the helical disruption (quantified by ALPHARMSD) for the S6 segment (S291-H299), reflecting the degree of local unwinding. b) Distribution of the minimum pore distance between residues 370-375 of the OR chain and residues 474-496 of the opposing Orco chain.

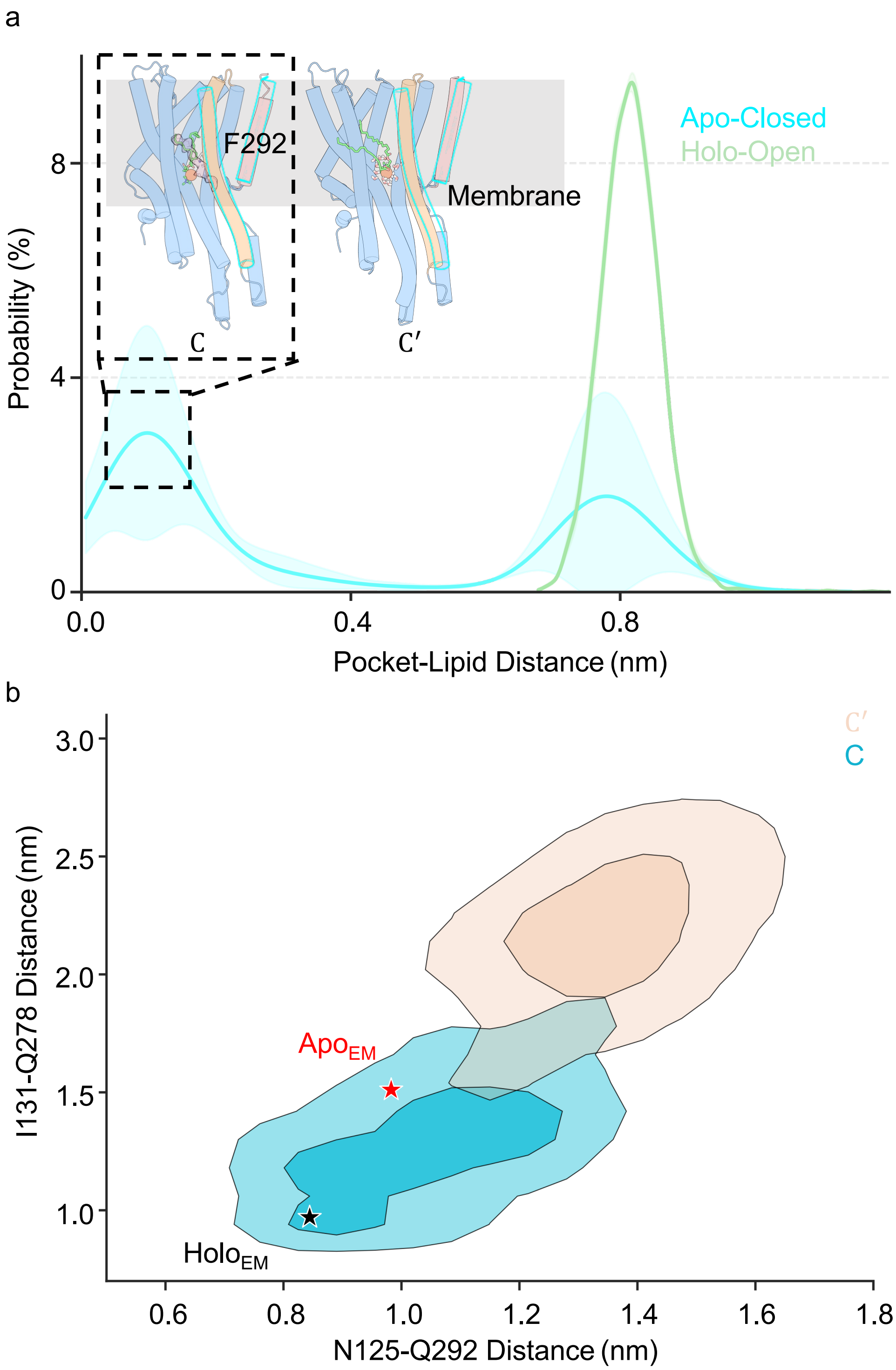


**Figure S11: Lipid insertion and S3-S6 geometry of the C′ state.** a) Minimum lipid-pocket distance distributions in the apo-closed and holo-open ensembles. The pocket center comprised OR residues 67, 68, 133, 136, 179, 180, and 183, which showed ligand-contact frequencies above 0.6 in the holo-open ensemble. Lines show means and shading indicates SE across three trajectories. Insets show representative C and C′ conformations sampled with POPC insertion. POPC is green, its phosphate headgroup is orange, the cryo-EM density is light purple, and water molecules are red and white. b) MSM-derived joint distributions of two distances across the upper S3-S6 interface in C and C′: I141-Q278 at its upper end and N125-Q292 at its lower end. Red and black stars indicate the apo-closed and holo-open cryo-EM structures, respectively.

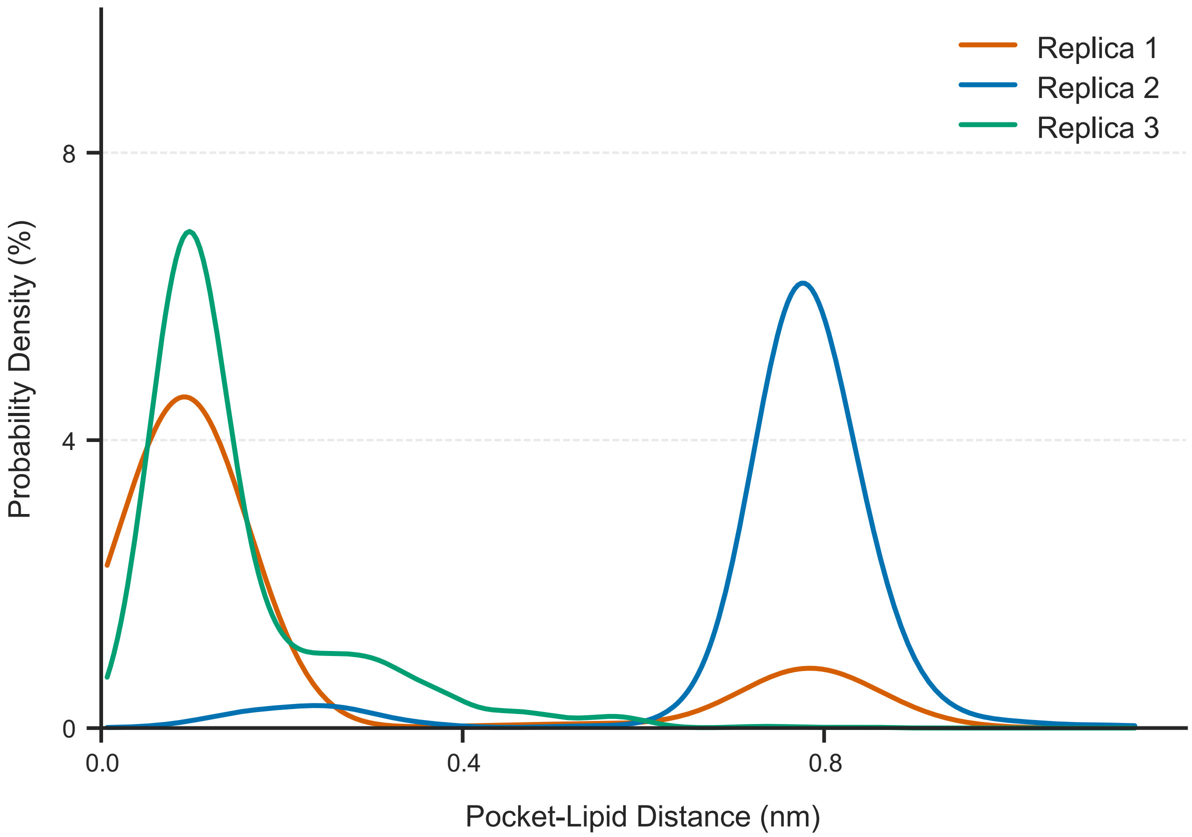


**Figure S12: Replica-resolved lipid-pocket distance distributions in the apo-closed unbiased simulations.** Probability-density distributions of the minimum distance between the lipid and the binding-pocket center for the three independent apo-closed replicas.

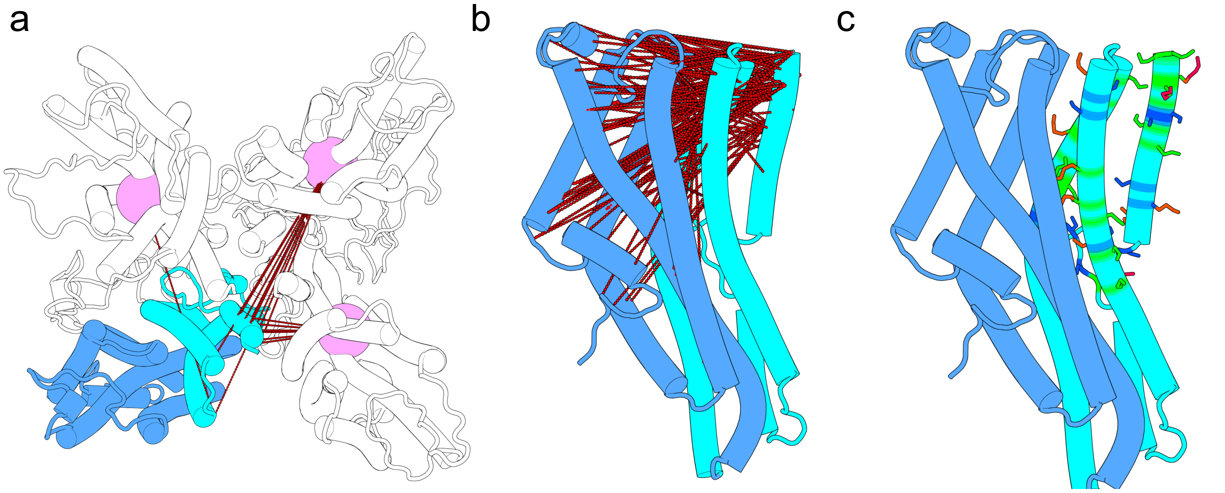


**Figure S13: The features ultimately input into the MSM construction.** a) Inter-subunit distances. b) Intra-subunit distances. c) Intra-subunit dihedrals; residues involved in dihedral features are displayed as sticks, with the specific dihedral angles colored as follows: $\chi_1$ in marine, $\chi_2$ in green, $\chi_3$ in orange, and $\chi_4$ in hotpink.
